# High-throughput selection of human *de novo*-emerged sORFs with high folding potential

**DOI:** 10.1101/2024.01.22.576604

**Authors:** Margaux Aubel, Filip Buchel, Brennen Heames, Alun Jones, Ondrej Honc, Erich Bornberg-Bauer, Klara Hlouchova

**Affiliations:** Institute for Evolution and Biodiversity, University of Muenster, Germany; Department of Protein Evolution, Max Planck-Institute for Biology Tuebingen, Germany; Department of Cell Biology, Faculty of Science, Charles University, Prague, Czech Republic; Department of Biochemistry, Faculty of Science, Charles University, Prague, Czech Republic; Institute of Organic Chemistry and Biochemistry, Czech Academy of Sciences, Prague, Czech Republic; Imaging Methods Core Facility, BIOCEV, Prague, Czech Republic

## Abstract

*De novo* genes emerge from previously non-coding stretches of the genome. Their encoded *de novo* proteins are generally expected to be similar to random sequences and, accordingly, with no stable tertiary fold and high predicted disorder. However, structural properties of *de novo* proteins and whether they differ during the stages of emergence and fixation have not been studied in depth and rely heavily on predictions. Here we generated a library of short human putative *de novo* proteins of varying lengths and ages and sorted the candidates according to their structural compactness and disorder propensity. Using Förster resonance energy transfer (FRET) combined with Fluorescence-activated cell sorting (FACS) we were able to screen the library for most compact protein structures, as well as most elongated and flexible structures. Compact *de novo* proteins are on average slightly shorter and contain lower predicted disorder than less compact ones. The predicted structures for most and least compact *de novo* proteins correspond to expectations in that they contain more secondary structure content or higher disorder content, respectively. Our experiments indicate that older *de novo* proteins have higher compactness and structural propensity compared to young ones. We discuss possible evolutionary scenarios and their implications underlying the age-dependencies of compactness and structural content of putative *de novo* proteins.

## Introduction

*De novo* protein emergence provides the genome with great innovative potential to explore the hitherto unexplored sequence space (McLysaght and Hurst 2016; Tautz and Domazet-Lošo 2011; Weisman and Eddy 2017; Van Oss and Carvunis 2019; Rödelsperger et al. 2019). In comparison to other known mechanisms of protein emergence that rely on recycling of conserved genetic elements, like duplication (Ohno 1970; Sikosek and Bornberg-Bauer 2010) or gene fusion (Dohmen et al. 2020), *de novo* genes emerge from non-coding regions of the genome (Bornberg-Bauer et al. 2021). *De novo* proteins are shorter on average than new proteins emerging through duplication (Montañés et al. 2023) and have low expression that is often restricted to specific tissues or conditions (Heames, J. Schmitz, et al. 2020; Wu and Knudson 2018; J. F. Schmitz, Chain, et al. 2020). Therefore, many potential *de novo* proteins are overlooked by classical annotation methods focusing on longer and highly expressed proteins. Due to the same challenges, short open reading frames (sORFs) and their microproteins remain understudied. Recently sORFs and microproteins have gained attention (Pueyo et al. 2016) and are proposed to serve as a reservoir for *de novo* proteins (Sandmann et al. 2023; Vakirlis et al. 2022). Additional *de novo* proteins are continuously detected across many species, including, e.g., several plants (Zhang et al. 2019; Marsch-Martínez et al. 2022), fruit flies (Heames, J. Schmitz, et al. 2020) and humans (Guerzoni and McLysaght 2016; Sandmann et al. 2023). Many of these studies focus on detection and functional characterisation of the *de novo* proteins, but few report structural characterisation (Lange et al. 2020; Bungard et al. 2017). Computationally, *de novo* proteins are mostly predicted to contain high structural disorder (Aubel et al. 2023; Dowling et al. 2020; Wilson et al. 2017; Peng and Zhao 2023). They may assume molten globule like structures containing secondary structure elements but lacking the stable tertiary fold of a globular protein (Bungard et al. 2017; Lange et al. 2020). Similarly, random-sequence proteins have been shown to contain secondary structure elements, but are best tolerated *in vivo* when they have a higher amount of disordered regions (Tretyachenko et al. 2017).

In a previous study comparing *de novo* protein candidates to random-sequence proteins (Heames, Buchel, et al. 2023) we showed experimentally that both sets of proteins are on average highly similar to each other concerning their solubility, interaction with chaperones and protease resistance. However, the putative *de novo* proteins showed slightly higher solubility, yet at the same time higher degradability when exposed to a bacterial Lon protease (Niwa et al. 2019) than their random-sequence counterparts. Their higher solubility combined with more degradation by the protease points to overall higher disorder content of *de novo* proteins compared to the random-sequence proteins. Corresponding to the experimental findings on both *de novo* and random proteins, solubility and prevention against aggregation seem to be the main bottleneck for newly emerging proteins to avoid purging by natural selection (Ángyán et al. 2012; Monti et al. 2021; Vakirlis et al. 2022; Agozzino and Dill 2018).

Here, we aim to select candidate *de novo* proteins originating from sORFs with high potential for compactness, and accordingly with a lower amount of disorder and increasing potential for folding. Our goal is to investigate whether and how frequently compact sORFs have the propensity to form secondary structure elements and potentially stable folds. We apply a high-throughput assay based on fluorescent activated cell sorting (FACS) of *Escherichia coli* cells to select top candidates from a library of 3750 putative *de novo* proteins (**Figure 1 a-b**). The design of the assay, based on previous work by **philipps°fret-based°2003**, makes use of Förster resonance energy transfer (FRET) between two fluorescent proteins with spectral overlap (Förster 1948). Efficiency of the transfer is inversely dependent on the distance of the fluorescent proteins, thereby offering a way to measure proximity of two molecules or to study intramolecular conformation states. By making use of the latter, we aimed to develop a high-throughput assay capable of screening for compact protein variants. In this assay, the library target protein is expressed in fusion with the FRET pair, serving as a linker between the fluorophores. The yellow fluorescent protein (YFP) serving as an acceptor is fused to the C-terminus of the target protein, while the donor cyan fluorescent protein (CFP) is at the N-terminus of the target protein (**Figure 1 c**). In case the two fluorescent proteins are connected by a stable and compact target protein, which places them within the FRET radius, the energy transfer can happen. Increased FRET signal reflects N- to C-terminal distance or persistence length of the linker proteins and can be interpreted as a measure of compactness (Rosmalen et al. 2017; Krishna and Englander 2005). Target proteins that are more flexible and do not provide a structurally stable link between the two fluorescent proteins, result in *E. coli* cells without FRET signal. The cells expressing the target protein with fluorescent proteins can be sorted accordingly using FACS. Thereby, we can select single protein sequences that have a low N- to C-terminal distance, are more compact and less disordered (**Figure 1 d**).

**Figure 1:**
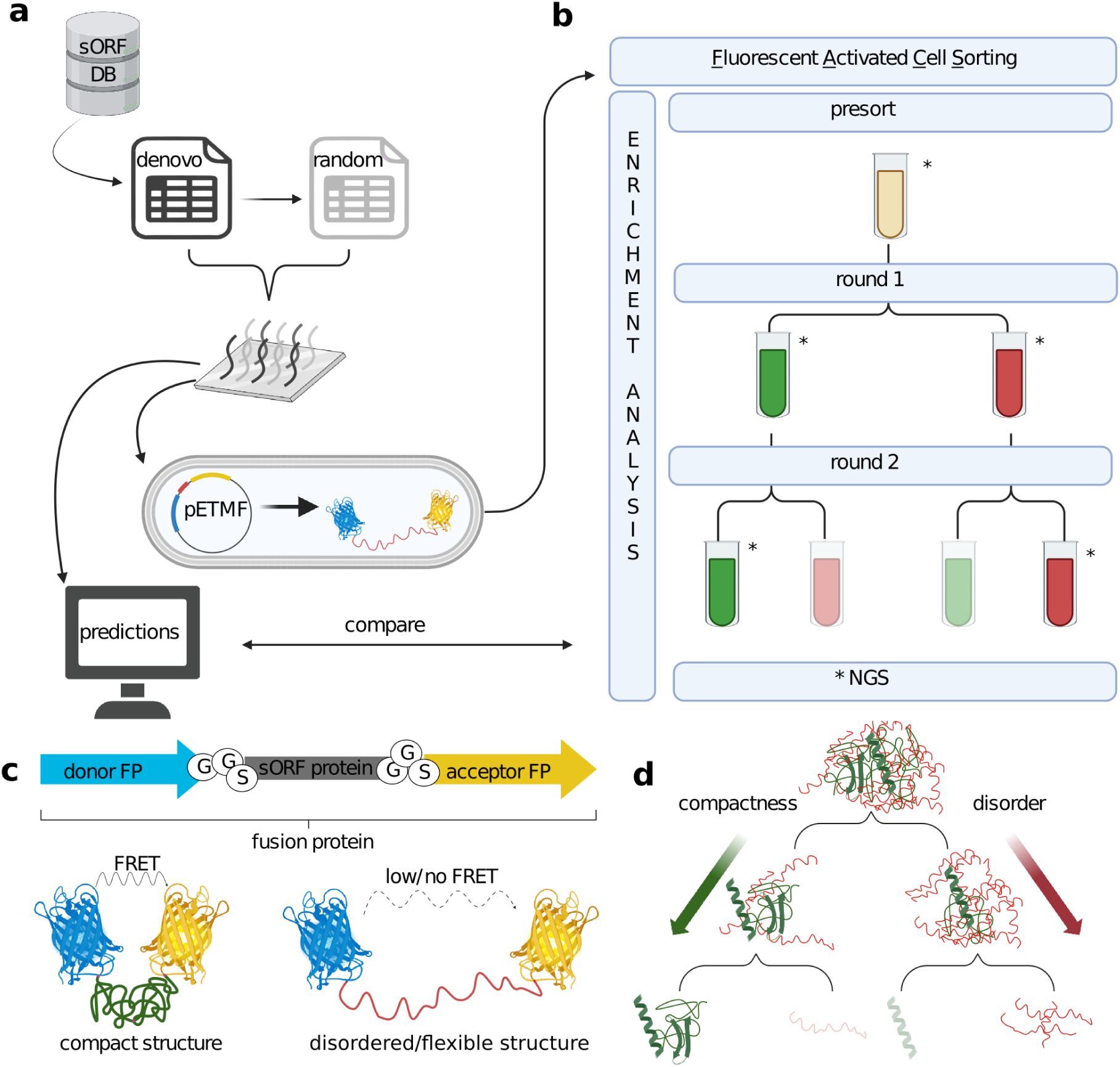
Workflow: a) We selected 3750 *de novo* emerged human sequences from the sORFs database and generated a library of comparable random sequences. Protein structure properties were predicted computationally. After ordering the libraries as oligonucleotides they were cloned into the pETMF plasmid and transformed into *E. coli* for the FRET-FACS assay. b) The FACS was performed in two sequential rounds for both libraries separately. Presorted cells containing single library sequences were sorted into FRET-positive (+, green) and FRET-negative (-, red) samples. Samples with * were recovered, sent for next-generation sequencing (NGS) and used for enrichment analysis. c) The library proteins (grey) are tagged with YFP (yellow) and CFP (blue) on the termini with GGS spacers. Compact library sORF proteins place the fluorescent proteins in close proximity and are expected to cause FRET. Disordered or fibrillar library proteins are expected to be FRET-negative. d) The presorted samples contain all library protein structures, the first round is expected to result in separation of compact and disordered structures, which becomes more pronounced in the second round. The FRET-negative proteins should have an increased N- to C-terminal distance and disorder, while the FRET-positive proteins show increased compactness and folding. Made with BioRender.

## Results

The library of 3750 putative *de novo* sORF proteins (DN) and corresponding random-sequence proteins (R) ranging from 32 - 59 amino acids were amplified from an oligonucleotide library synthesis (OLS) pool. DN and R libraries were separated in PCR by using library-specific primers and then sequenced using Illumina next generation sequencing (NGS). The PCR product of both libraries covered 90 % (3365 DN and 3367 R) of the initial libraries (**Figure S1**). Library sequences of DN and R were then cloned into pETMF plasmid, placing the single library sORF proteins as a linker between two fluorescent proteins. Upon expression in *E. coli* the library proteins are tagged with CFP at the N-terminus and YFP at the C-terminus resulting in a fusion protein (**Figure 1 c**). More compact linker proteins lead to a stronger FRET signal between the two fluorescent tags and can be sorted using FACS on the *E. coli* expressing the fusion protein. Each library was sorted into FRET-positive (FRET+) and FRET-negative (FRET-) categories based on controls (GS-linkers and previously characterised proteins of the same length, see **Table 1**) in two sequential rounds starting from the presorted samples. After completing all FACS rounds, cells from the different samples were recovered separately, barcoded and their DNA sequenced using NGS.

### Fluorescence Lifetime measurements

We chose mTurquiose2 and mVenus as a FRET pair, as both of these fluorescent proteins express well in *E.coli*, possess self-association preventing mutations, exhibit high brightness and, in the case of mTurquiose2, long fluorescence lifetime (Bajar et al. 2016). First, we generated a cassette consisting of the FRET pair genes separated by a Golden Gate assembly compatible cloning site and inserted the cassette into the backbone of the pET24a(+) vector. The resulting vector pETMF allows for T7 promoter controlled expression of the large fusion protein, where the protein of interest is connected to the FRET pair by one glycine-serine repeat and glutamate and leucine residues as part of the cloning process. To validate the performance of our compactness assay, we selected several well characterised *E.coli* proteins with different properties and generated constructs with the FRET pair separated by GGS repeat linkers of various lengths (**Table 1**). Intensity based FRET efficiency measurements require purified samples of known concentrations. Therefore we used time correlated single photon counting (TCSPC) to obtain a fluorescence lifetime of the donor molecule, which can be used to calculate FRET efficiency *in vivo*. Fluorescence lifetime is a property inherent to every fluorescent molecule and it defines the time it takes the molecule to return to its ground state upon excitation. In the presence of a FRET acceptor, the lifetime of the donor shortens and by comparing it to the lifetime of the donor alone the FRET efficiency can be calculated. After exponential curve fitting, we deconvoluted the fluorescence lifetime of the donor alone and fusion protein constructs. The fluorescence lifetime of the donor, 3.79 nanoseconds, slightly deviates from lifetime of 4.0 nanoseconds reported in a previous study (Goedhart et al. 2012), which can be attributed to fluorescence quenching of the cellular environment in bacteria. Nevertheless, we observed an increase in lifetime with increasing length of the linker in case of GS-linker controls. The various measured lifetimes for selected *E.coli* proteins corresponded to their overall flexibility and structure content, albeit invariant of the length (**Table 1**).

**Table 1:** Fluorescence lifetime of different control proteins measured *in vivo*. GS2/3/4 - controls with corresponding number of GGSGGS repeats; 1259 and 665 - characterised random proteins (Tretyachenko et al. 2017); BolA - transcriptional regulator; IscA - iron binding protein, forms homodimers around Fe atoms; SodA - superoxide dismutase; CspA - cold shock DNA binding protein; FtsB - transmembrane protein of bacterial divisome; YacG - DNA gyrase inhibitor. FRET efficiency is calculated as: 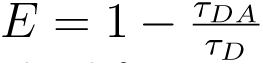, where τ_DA_ is the lifetime of the donor in presence of the acceptor and τ_D_ is the lifetime of the donor alone.

| Protein name | Length (aa) | Lifetime measured (ns) | FRET efficiency | Properties |
| --- | --- | --- | --- | --- |
| mTurquoise2 | N/A | 3.79 | N/A | Donor |
| GS2 | 12 | 2.51 | 0.338 | Shortest linker |
| GS3 | 18 | 2.85 | 0.248 | Short linker |
| GS4 | 24 | 2.9 | 0.235 | Medium length linker |
| 1259 | 100 | 3.36 | 0.113 | Disordered |
| 665 | 100 | 2.64 | 0.303 | Secondary structure rich |
| BolA | 107 | 2.56 | 0.325 | Globular, terminal disorder |
| IscA | 107 | 2.76 | 0.272 | Homodimeric |
| SodA | 206 | 2.77 | 0.269 | Homodimeric |
| CspA | 69 | 2.8 | 0.261 | β-barrel |
| FtsB | 103 | 3.41 | 0.1 | Coiled coil |
| YacG | 68 | 3.56 | 0.061 | Disordered |

### Flow cytometry

Following the fluorescence lifetime measurements, continued with flow cytometry experiments to verify that spectroscopic results correspond to the cytometric ones. The nature of our assay, being an intramolecular FRET sensor, does not require the tedious control process as in the case of detecting protein-protein interactions by FRET (Banning et al. 2010). Nevertheless, we employed 3 channels - donor excitation and emission (donor channel), acceptor excitation and emission (acceptor channel), donor excitation and acceptor emission (FRET channel). First, we used donor and acceptor channels to gate a population positive for both fluorophores (P1), suggesting presence of a complete fusion protein (**Figure S4**). To robustly detect the FRET signal, we derived a parameter of ratio between the FRET channel and the donor channel (FRET ratio). We observed that the FRET ratio of the control proteins were in line with the FRET efficiencies obtained from fluorescence lifetime measurements (**Figure S4**). We then proceeded to sort the DN and R library samples. First, we gated approximately 30 % of the events exhibiting maximum fluorescence for the donor and the acceptor channel (P1 in **Figure 2 a**). Apart from the main double fluorescent population, a smaller population showing only donor signal appeared, which can be attributed to spurious stop codons or variants difficult to translate. After projecting the P1 population to the FRET ratio histogram, we started with excluding the top 3 % of the population, as the cells showing extreme fluorescence have lower viability. FRET- and FRET+ gates were set by gating the bottom and the top 10 % of the distribution, with 30 000 events sorted for each of these gates (**Figure 2 a**, left and middle). Sorted cells were recovered on LB-agar plates and new expression cultures prepared (see Methods). In subsequent round of FACS, the FRET+ population was only sorted for high FRET signal and FRET- population for low FRET signal, to further strengthen the selection (**Figure 1**). Overlaid FRET ratio projections of P1 populations from two rounds of sorting (**Figure 2 a**, right) showed an incremental shift of the FRET ratio signal away from the naive population (DN all). This trend was most prominent in the FRET-negative samples, however in the second round of FRET-positive sorting the FRET ratio drops with the median even below the naive population. One possible explanation for this phenomena might be more structural variety of the selected variants, resulting in larger spread of the signal. The predictions of all sorted sequences from round one to two follow the expected trend with decreasing disorder and increasing compactness and secondary structure content (**Figure S3**).

**Figure 2:**
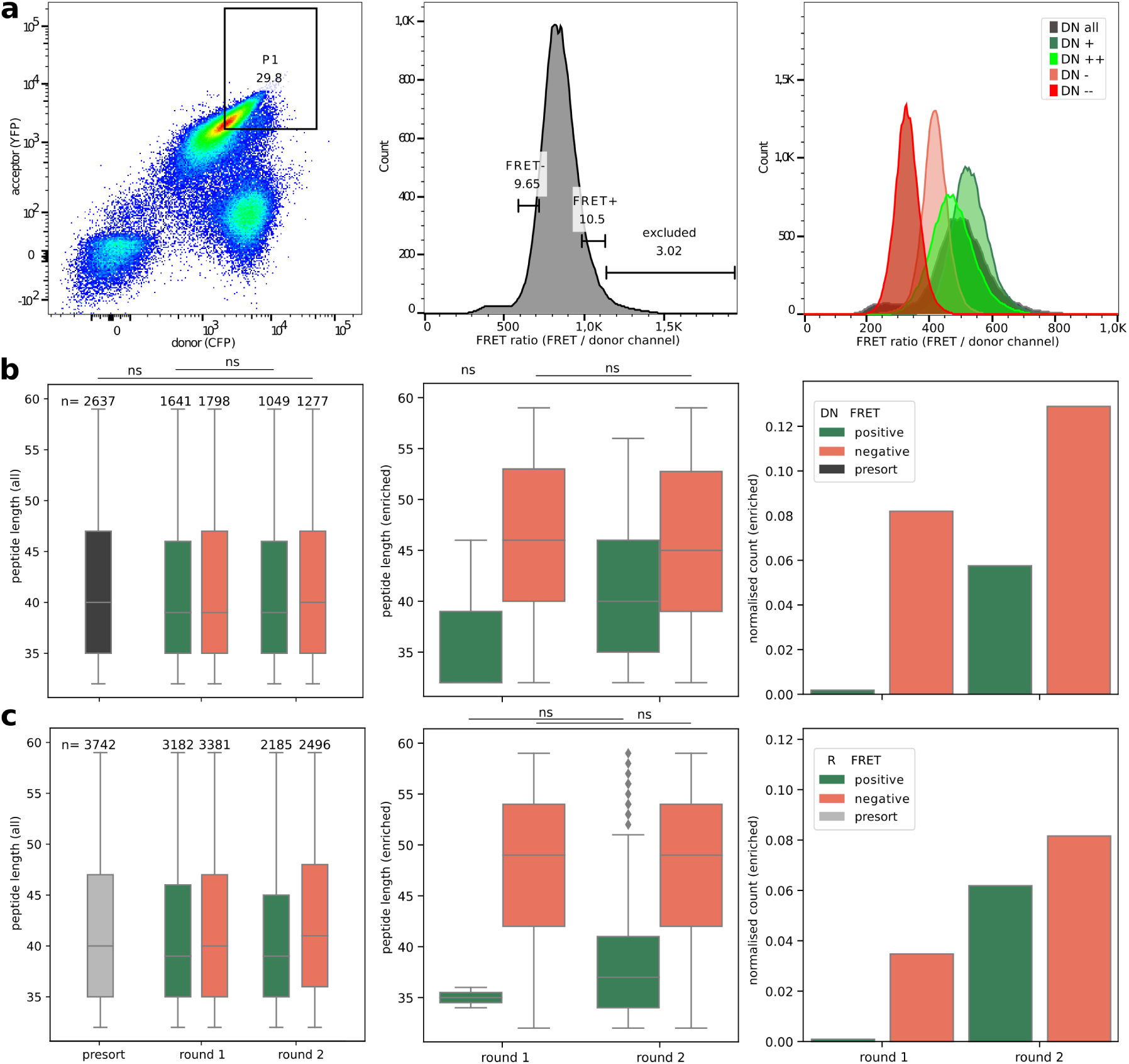
a) On the left, exemplary gating of the double fluorescence positive population P1 is shown. The central histogram shows the FRET ratio of the P1 population with arbitrary gate ‘excluded’ (see Results) and FRET-/FRET+ gates. The right plot shows the FRET ratio of overlaid P1 populations after two rounds of sorting, recorded after the sorting experiment on LSRFortessa cytometer. b, c) Peptide lengths of all sequences of libraries DN (b) and R (c) in FRET-positive and FRET-negative samples across rounds including total number of unique sequences on top of the bars (left), peptide lengths of significantly enriched sequences (centre) and number of enriched sequences normalised by total sequences present for each library (right). The lengths are significantly different (p-value ¡ 0.05) according to Tukey HSD test between all categories if not indicated otherwise (ns).

### Sorting of DN and R libraries

Over two thirds of the presorted sORF proteins from the DN and R libraries were present at least once in the FRET-positive and FRET-negative samples of round one, indicating that a second round of sorting is needed to clearly separate the FRET-positive from FRET-negative proteins (**Figure 2 b-c**, left). After the second round of sorting, the number of sORF proteins decreased to around half of the presorted sORF proteins in FRET-positive and FRET-negative samples for the DN library. In contrast, two thirds of the presorted sORF proteins of the R library are still present in both samples after the second sorting. There is no significant difference in length between FRET-positive and FRET-negative sorted sORF proteins after the first round of sorting for both DN and R libraries. After the second sorting both libraries contain significantly shorter sequences on average in the FRET-positive samples than the FRET-negative samples. However, the whole length range of library sequences is covered in both FRET-positive and FRET-negative samples.

### Enrichment analysis

We calculated enrichment of single sequences in FRET-positive and FRET-negative samples between the rounds to determine the sORF proteins that are over represented in the different FRET categories. Overall, more sequences are significantly enriched (p-value *<* 0.05) in the FRET-negative sample compared to the FRET-positive for both DN (**Figure 2 b**) and R (**Figure 2 c**). A higher proportion of the presorted library is enriched in the FRET-negative sorting in the DN library compared to R (0.12 and 0.08, respectively, after round two of sorting). The proportion of enriched sequences increases substantially from close to zero in the first round of sorting to 0.06 in the second round for both DN and R. Comparisons between FRET conditions of enriched proteins in round one are difficult because of very few significantly enriched sequences in the FRET-positive samples. From round one to two, the average length of the FRET-positive enriched proteins increases while the predicted disorder decreases. This increasing length could be causing the drop in FRET ratio observed for DN after the second round of sorting (see **Figure 2 a**). The FRET-negative samples do not change in length, but predicted disorder increases from round one to two for both libraries (significant for R). Four sORF proteins of the DN library are enriched in the FRET-positive samples as well as in the FRET-negative samples, and are discarded as false positives when choosing most compact candidates with folding potential.

For a comparison of protein properties between FRET-positive and FRET-negative samples, we took only the proteins that are significantly enriched from presort to round two. This way we use the highest number of sORF proteins and the most confidently enriched ones (see **Figure 2**). As the predictions for random-sequence proteins are less reliable (Middendorf and Eicholt 2024), from hereon only the DN library is regarded for analysis (for library R see supplementary **Figure S5, S8,S9**). The protein sequences enriched in the FRET-negative samples are significantly longer on average than sequences that are enriched in the FRET-positive samples. To predict structural properties, we used ESMFold (Lin et al. 2023), which has been proposed to be more reliable on sequences without homology (Elofsson 2023), like *de novo* proteins. Taking the ESMFold predictions, we calculated the percentage of secondary structure (helix, sheet, coil, PP-II helix, turns), radius of gyration, average solvent accessibility per amino acid (asa) and N- to C-terminal distance. Sequences enriched in FRET-positive are predicted to have significantly lower radius of gyration, lower Nto C-terminal distance, lower average solvent accessibility, higher amount of secondary structure elements, and higher confidence of ESMFold predictions seen by the higher pLDDT (**Figure 3** and **S5**).

**Figure 3:**
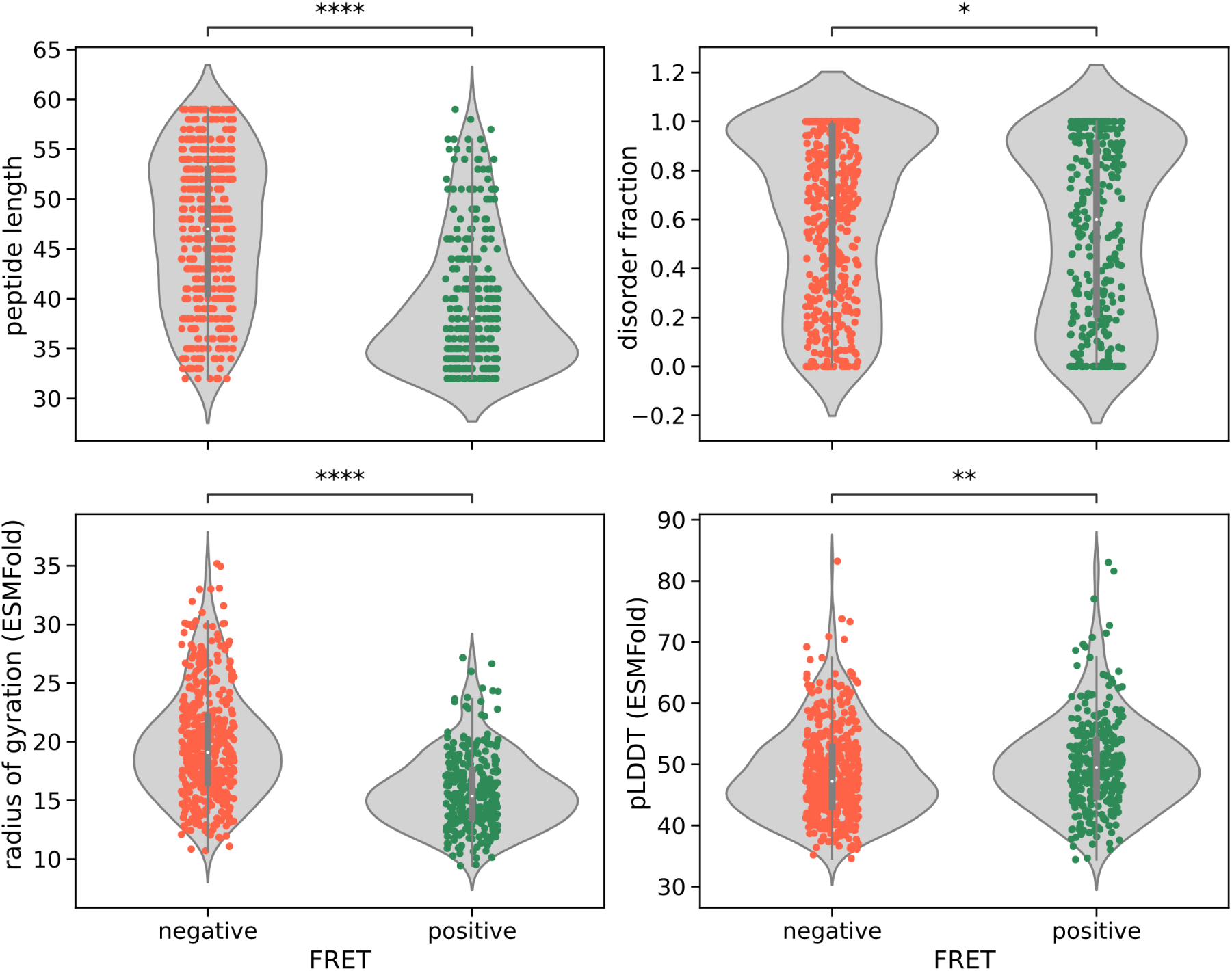
Protein property predictions (y-axis) of significantly enriched sequences in round two of sorting for FRET-positive (green) and FRET-negative (red) subsets of the DN library. Stars indicate significance calculated with t-test (p-value *<* 0.05 * *<* 0.01 ** *<* 0.001 *** *<* 0.0001 ****).

### Predictive models

We used predictive modelling to gain further insights into which predicted protein properties had an impact on whether *de novo* sORF proteins are sorted into the FRET-positive or FRET-negative samples. For this purpose we split all enriched sequences into test and training data sets and used elastic net type regression (Tay et al. 2023) of the log fold change (LFC) from presorted samples to round two of sorting plotted against different protein properties (**Figures 4, S7, S6**). The protein properties with the biggest influence on FRET-positive sorting are predicted turns in the structure and disorder, while length of the protein sequence has only a very low impact (**Figure 4**). The LFC is positively correlated with predicted turns after secondary structure elements, predicted beta-sheets and age of transcription but negatively correlated with length, predicted disorder and coils, radius of gyration and N- to C-terminal distance (**Figure S6**). The length of the sORF proteins had a bigger impact (correlation 4x higher) on whether a protein is enriched in the FRET-negative samples compared to FRET-positive. Predicted disorder and age categories (transcription or BLAST) have the reverse effect on FRET-negative sorting compared to FRET-positive sorting, but of the same magnitude (-2.2 vs 2.6 and 0.06 vs -0.06).

**Figure 4:**
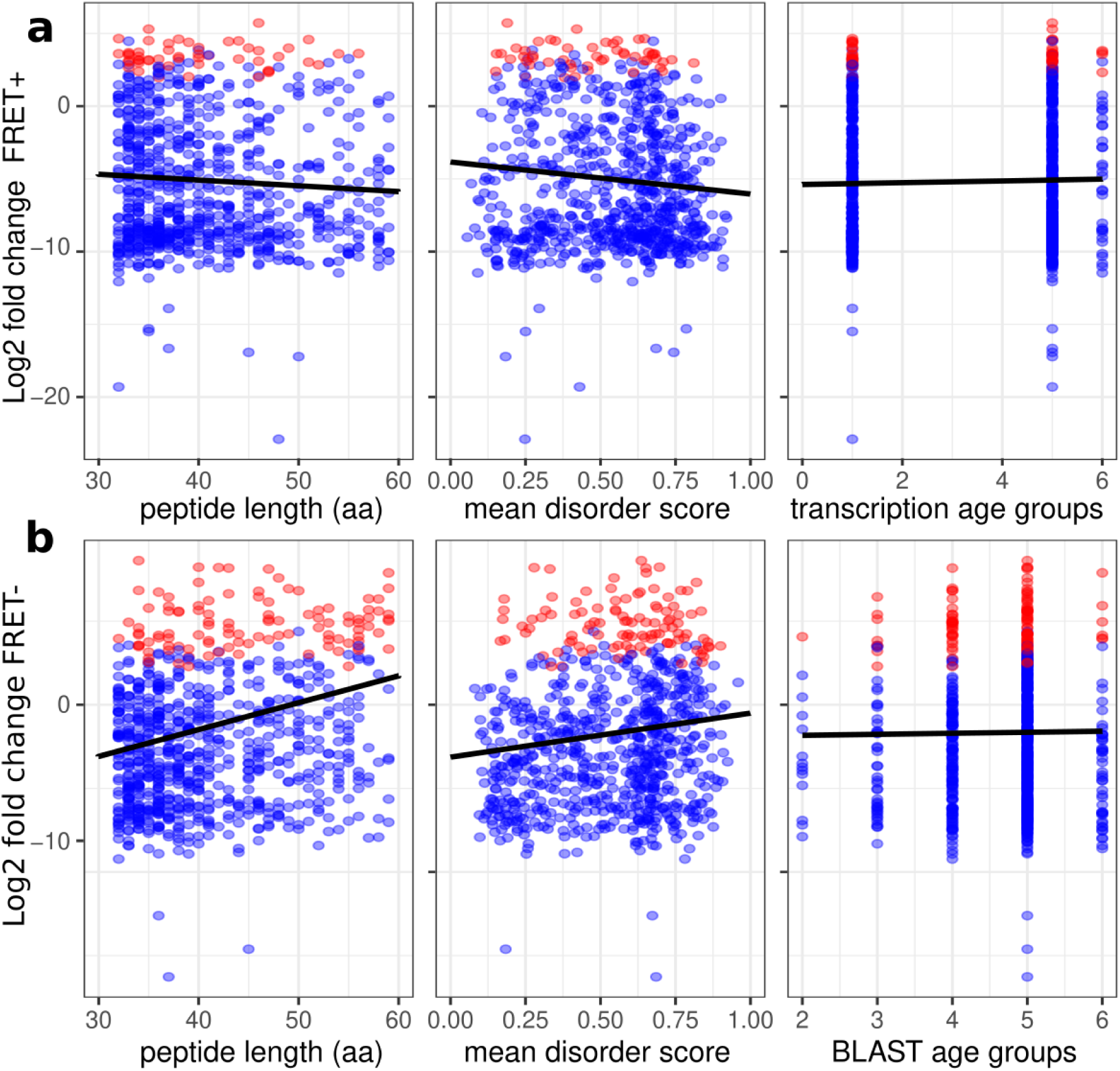
Predictive plots for FRET-positive (a) and FRET-negative (b) *de novo* sORF proteins enriched from presort to round two. The points are the test data set with significantly enriched sequences in red. The line is the coefficient of the particular predictor with all other variables either set to zero or to the median value. Note that in an an elastic net type regression as used here, the uncertainty is not calculable in the same way as a normal regression, so no confidence intervals are added.

### Ages of *de novo* sORFs

Protein folding and compactness is usually considered to be a trait that needs long evolutionary timescales to develop, given that it is highly unlikely for a random sequence. Young *de novo* proteins and random-sequence proteins are therefore predicted to be highly disordered, lacking secondary structure elements and a stable tertiary fold. A major unresolved issue in understanding the evolution of *de novo* proteins is, whether older *de novo* proteins are more structured than younger ones. Our DN library contains putative *de novo* emerged sORFs of different ages. We grouped sequences into age categories 1-6, covering an evolutionary age range from 6-90 Million years (My). The age categories are based on: i) BLAST age, i.e., significant sequence similarity in outgroup species (chimpanzee, gorilla, orangutan, macaque and mouse) or ii) RNA age, i.e., detectable transcription in human, macaque and mouse. There are no sequences of blast age 1 (human only), because we excluded sequences without any sequence similarity in related species in the library design as the mechanism of emergence could not be determined. The *de novo* sORFs specific to monkeys (ages 2-5) show similar behaviour in the assay, with around 70 % of sequences in the FRET negative enriched, i.e., disordered proteins and 30 % in the FRET positive enriched, i.e., more compact sORF proteins (**Figure 5**). The oldest age group with homologous sequences detectable up to mouse (age 6) has a higher percentage of FRET positive enriched sequences ( 45%) than the younger sORFs. For the age groups based on transcription, we see a similar trend. The older *de novo* sORFs that are transcribed in macaque (age 5) or mouse (age 6) are more frequently represented in the FRET positive enriched sequences than the younger ones which are transcribed in human only (age 1). The predictive models described above also predict a positive correlation between LFC and age of transcription for the sORF proteins enriched in the FRET-positive samples (**Figure 4**). The observed trend deserves further verification using larger data sets and other species because only few enriched sequences (n=12 and n=18) belong to the oldest age group 6 and differences between age groups are not statistically significant (p-value ¿ 0.05, chi2 test).

**Figure 5:**
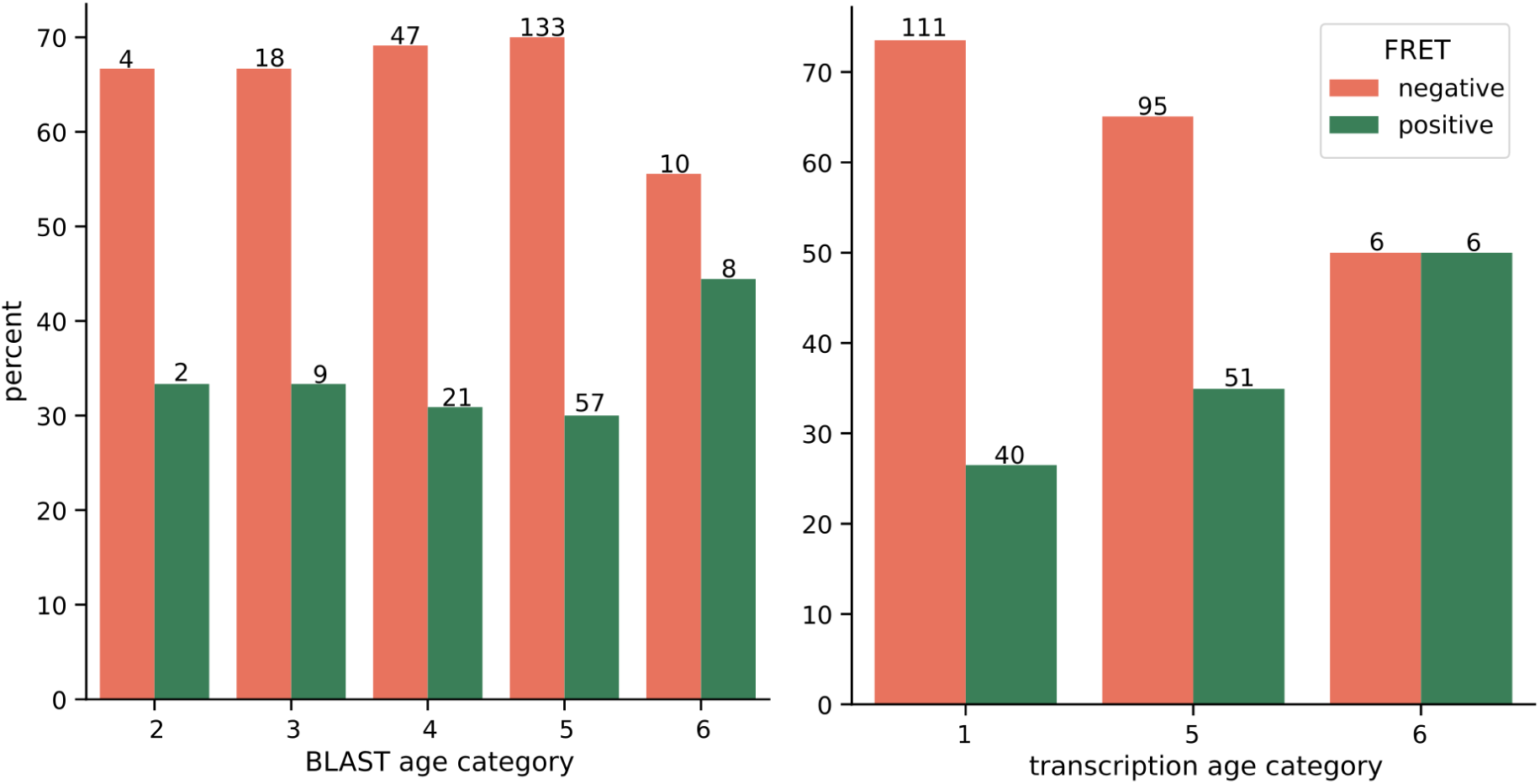
Percentage of all significantly enriched sequences in the DN library separated by FRET-positive and FRET-negative enrichment sorted by BLAST age categories (left) and transcription age categories (right).

### Top enriched *de novo* sORFs

The aim of this study was selection of putative *de novo* sORF proteins with the potential ability to fold into more compact structures. To verify the selection of appropriate candidates from the library of 3750 sORF sequences, we used the ESMFold predictions of sequences with highest enrichment in the FRET-positive and FRET-negative samples after two rounds of sorting (**Figure 6**). As an additional filter for FRET-positive sORF proteins, we took the candidates enriched in the first round as well as the second round, considering the drop in FRET ratio in the second rround of sorting (**Figure2 A-C**). The top enriched FRET-positive structures (A-D) contain a comparably high amount of predicted *α*-helices and display a relatively compact structure. The radius of gyration for the top structures is below the average, as calculated for the DN library based on ESMFold predictions. The top FRET-negative enriched sORF proteins (E-G) are predicted to consist of mainly disordered regions, with only one out four structures having a small *α*-helix predicted. The confidence score (pLDDT) of predicted structures for the top enriched sORFs is low, within the range of the average for all *de novo* sORF library proteins. Although we observe the trend for more secondary structure and less disorder for FRET-positive sequences, a few FRET-positive sORF proteins still show high amounts of predicted disorder and low secondary structure content underlining the importance for additional filtering in future studies.

**Figure 6:**
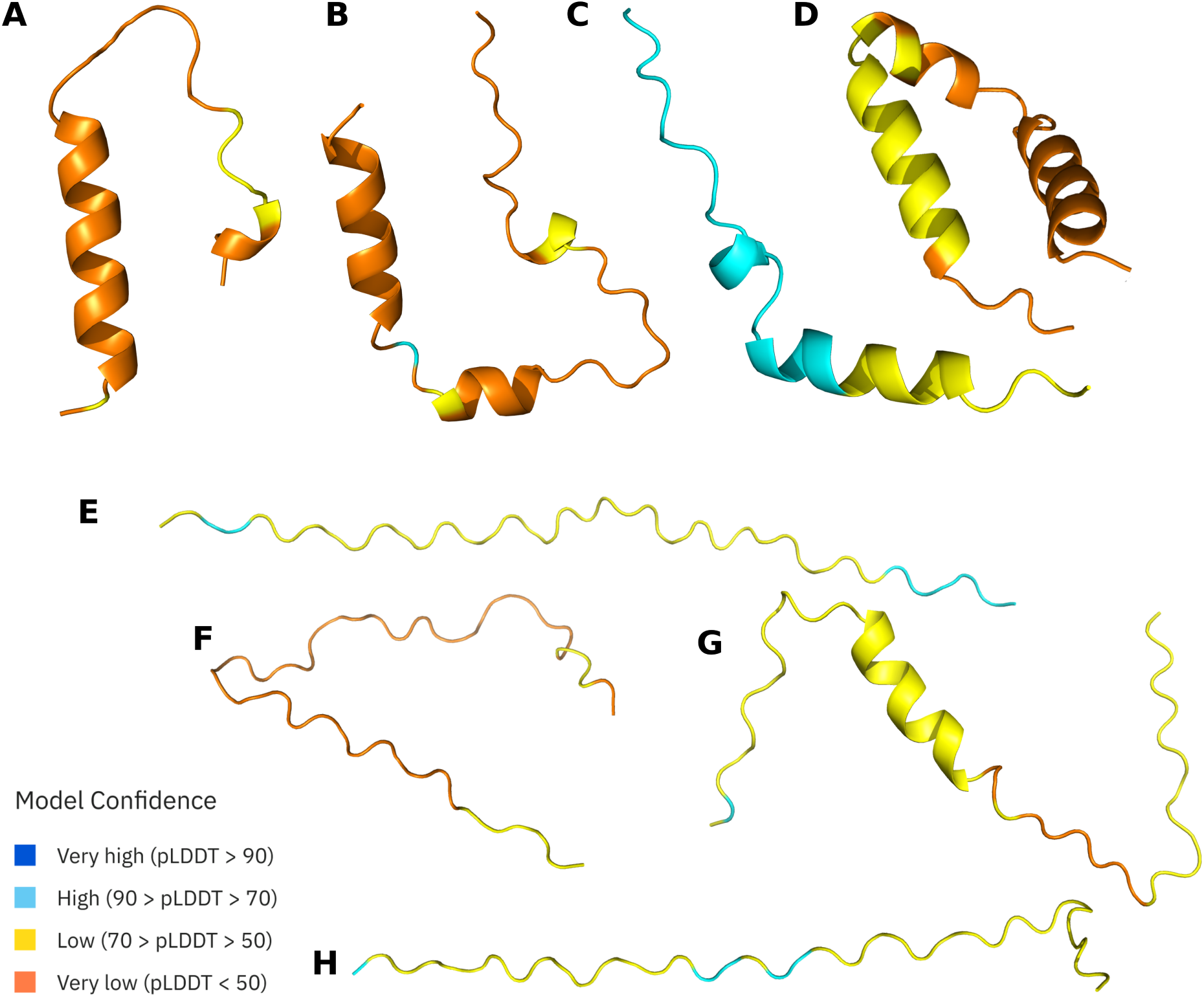
Examples for predicted *de novo* sORF protein structures of the four most confidently enriched FRET-positive (A-D) and FRET-negative (E-H) sequences coloured by model confidence.

While the top enriched sORF proteins correspond to the overall expected predictions, they do not represent the best candidates according to the predicted protein properties, i.e., with the lowest radius of gyration and highest amount of secondary structure. All top enriched proteins, FRET-positive and FRET-negative, are indistinguishable from other enriched proteins according to predictions. However, looking at the predicted disorder in relation to length (**Figure 7**), the FRET-positive enriched sORF proteins are scattered mostly at the lower edge of the plot with decreasing disorder for longer sORFs. The FRET-negative enriched sORF proteins do not display a trend in the distribution.

**Figure 7:**
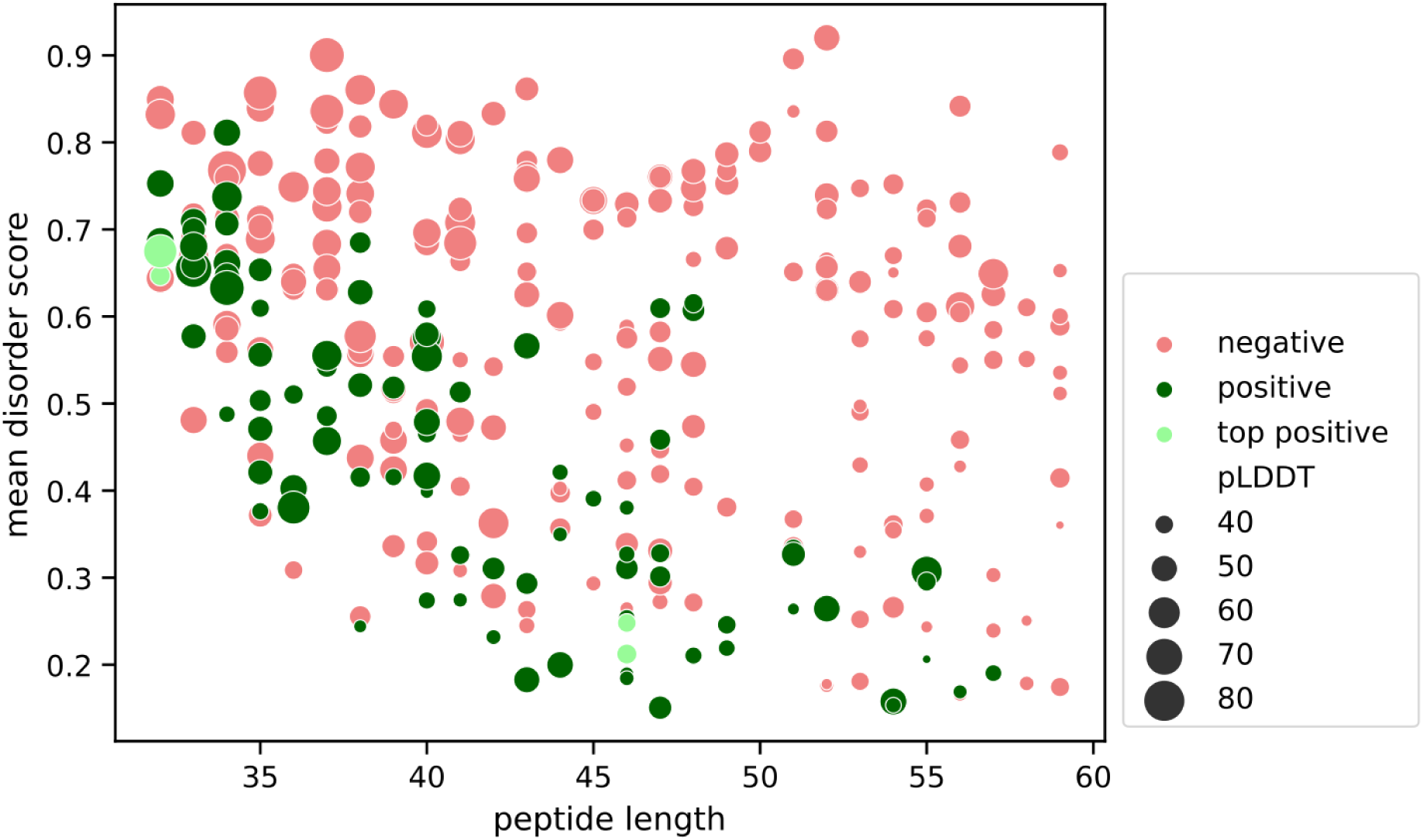
Scatterplot of significantly enriched sORF proteins looking at predicted mean disorder and length. Each dot represents a single sORF protein, the size of the dots corresponds to the confidence measure pLDDT of ESMFold predictions.

## Discussion

In this study, we screened putative *de novo* sORF proteins of different ages for candidate sequences with high compactness and with that, the capability for folding. Using an assay that combines FRET between an N-terminal YFP and a C-terminal CFP of the target protein with FACS allows for High-Throughput screening of thousands of candidate sequences. We expected the FRET-positive sORF proteins to have a low N- to C-terminal distance, resulting in a higher probability for compactness and folding. The FRET-negative sORF proteins, on the other hand, are expected to have a higher N- to C-terminal distance and higher amount of disorder in their structure. The candidate sORFs are of varying lengths (32-59 aa) to screen as many *de novo* candidate sequences as possible. Due to the distance-based nature of the assay, shorter sequences are more likely to trigger the FRET than longer sequences, and are therefore present in higher numbers in the FRET-positive samples. We still observed some of the longer sequences enriched in the FRET-positive sorted cells, especially after the second round of screening for the *de novo* library, which might explain the shift towards lower FRET ratio (**Figure 2 a**). According to predictive modelling, the length of sORF *de novo* proteins has little impact on sorting for the FRET-positive enrichment. For the FRET-negative enrichment, the length has a coefficient four times higher than for the FRET-positive sequences indicating a higher impact of length for the sorting of FRET-negative sequences. As expected, the length is correlated negatively with LFC for FRET-positive enriched sequences and positively for FRET-negative ones (**Figure 4** ). The disorder, as predicted, influences both FRET-positive and FRET-negative enrichment in the same order of magnitude but opposite signs as expected. For FRET-positive sequences, there is a negative correlation between LFC and predicted disorder while there is a positive correlation for FRET-negative sequences. In line with the literature on *de novo* proteins and random sequence proteins, a higher percentage of both libraries is FRET-negative and with that, predicted to be rich in disordered regions and lacking a stable tertiary fold (Heames, Buchel, et al. 2023; Bornberg-Bauer et al. 2021). In a previous study on *de novo* and random proteins (Heames, Buchel, et al. 2023) we observed that *de novo* proteins tend to be more soluble than random proteins of same lengths and amino acid frequency, mainly because random proteins have higher propensity for secondary structure. Here, we observe a similar trend with a slightly higher percentage of *de novo* sORF proteins enriched in the FRET-negative samples with high disorder predicted compared to the random ones (**Figure 2 b-c**). This can be explained by the evolutionary pressure on newly arising proteins to be soluble and to not disturb the function of the cell and to not cause harmful aggregation (Monti et al. 2021; Ángyán et al. 2012; Vakirlis et al. 2022; Agozzino and Dill 2018).

One of the remaining questions in *de novo* protein research is, if older *de novo* proteins are more structured and contain fewer disordered regions than newly emerging *de novo* proteins. A stable tertiary fold is difficult to attain from scratch (as in *de novo* emergence) and probably needs time to be formed by evolution (Bornberg-Bauer et al. 2021). Here we observed a trend for older *de novo* sORF proteins to be enriched in the FRET-positive samples compared to the younger ones (**Figure 4 - 5**). This trend might indicate a higher propensity for folding in the older *de novo* proteins as has been hypothesised previously (Wilson et al. 2017; J. Chen et al. 2023; Middendorf and Eicholt 2024). To best of our knowledge for the first time, we observe this trend experimentally, though further verification with a higher number of older and longer *de novo* proteins is needed. Overall, these results taken together with our earlier study (Heames, Buchel, et al. 2023), demonstrate how large libraries comprising proteins with random sequences or putative *de novo* proteins can be studied in a tractable experimental setup. While our results regarding structural properties are generally in good agreement with computational predictions, several outliers demonstrate that, wherever possible, experimental confirmations are recommended particularly for single protein studies (Terwilliger et al. 2023). Akin to earlier computational studies on age stratification of *de novo* proteins we find that older *de novo* proteins, i.e., those which have orthologs and transcription in several outgroup species, are generally more compact and have a higher propensity for folding (**Figure 4,5**). This may hint at an evolutionary process which favours *de novo* proteins that survive purging by drift to become longer and to assume a more foldable scaffold. Almost all *de novo* proteins are much shorter than the optimal globular protein domain of an average 165 aa (Shen et al. 2005), often containing fewer than 100 amino acids (Heames, J. Schmitz, et al. 2020; Blevins et al. 2021). Therefore, an optimal surface to volume ratio is difficult to accomplish with the hydrophobic-polar pattern along the peptide chain. To generate such a hydrophobic-polar pattern from a peptide chain which emerges from a DNA stretch which has most likely been subject to a strong and largely unconstrained drift for a long evolutionary time, positive selection would be necessary (Agozzino and Dill 2018; Shen et al. 2005). While many studies look for selection on *de novo* protein-encoding sequences (Zhang et al. 2019; Zhao et al. 2014; Heames, J. Schmitz, et al. 2020), only few studies so far have found convincing evidence for *de novo* proteins being under positive selection (Gubala et al. 2017). Furthermore, computational studies on the extension of ORFs by read-through and loss of stop codons (Kleppe and Bornberg-Bauer 2018; Klasberg et al. 2018) also suggested that extending the peptide chain from previously non-coding regions will result in a higher degree of disorder, at least in first place. Alternatively, it is conceivable that older *de novo* proteins are not or only marginally extended because they are ”born this way”, as suggested recently by Peng and Zhao (2023). This hypothesis would be supported by the loss dynamics of *de novo* proteins which suggests that *de novo* genes emerge more often than duplicates but are lost much faster such that, after longer evolutionary time scales, the fraction of *de novo* emerged genes among novel genes has become much smaller (J. F. Schmitz, Ullrich, et al. 2018; Grandchamp et al. 2023; Montañés et al. 2023). Our data do not allow for differentiating between these scenarios, but our results lay a foundation for future studies.

## Methods

### Identification of *de novo* sORFs

Human sORFs (n=2,626,006) were downloaded from http://sorfs.org. sORFs less than 30 amino acids long were discarded and the longest isoforms selected. The resulting set of 53,670 sORFs were used as a query against the NCBI-nr database. diamond was used with arguments: blastp -p 28 -e 0.001 --sensitive -b 2.75. sORFs with significant hits (threshold 1e-3) in a genus other than *Homo* were discarded as non human-specific. The apparent emergence mechanism of the remaining (human-specific) orphan sORFs was subsequently obtained by mapping of each ORF to the outgroup genomes of four primate genomes as well as the *Mus musculus* genome (see table S1 for accession numbers). First, BLAT was used to map each ORF against the six-frame translations of all five genomes. BLAT hits were filtered to only include the best hit for each reference sequence using the script pslCDnaFilter with arguments -maxAligns=1. Subsequently, we took a conservative approach to find the highest ranking annotation that overlapped with the set of BLAT hits across all outgroups for each sORF. Only if all BLAT hits across all outgroup genomes did not overlap with any annotated features (as defined in the corresponding Ensembl gtf files) did we define a sORF as ‘intergenic *de novo*’. Alternately, if any of the BLAT hits overlapped with a gene feature but no CDS feature, we defined it as ‘intronic *de novo*’ or ‘UTR *de novo*’. All sORFs overlapping with any annotated CDS features were discarded as non *de novo* resulting in a total number of 6649 *de novo* sORFs.

### Library design and oligo specification

Two libraries, DN (*de novo*) and R (random) were designed *in silico*. We used the set of 6649 *de novo* sORFs identified (see ) as a starting point for library design. First, any homologous *de novo* sORFs were discarded using cd-hit (see script remove similar.py). Coding sequences (CDSs) for library R were then generated by random selection of amino acids using the frequency of amino acids in library DN, with sequence lengths also matched to those of library DN. Final oligonucleotides were specified by addition of upstream and downstream barcodes, allowing each library to be PCR amplified separately from the oligo pool. DnaChisel (Zulkower and Rosser 2020) was subsequently used to codon optimise CDS regions for protein expression in *E. coli*, while avoiding introduction of unwanted restriction sites (inc. BsaI). Codon optimisation of the target ORF by selecting the best target species codon possible, given additional optimisation constraints. These two subpools of 3750 sequences were specified within a pool of 7500 oligos in total. For each ORF, flanking regions encoding BsaI sites and overhangs were added, and start codons were replaced with the canonical ATG start if it was not already present. Primers (16 bp) unique to each subpool were then added, and if the oligo length was less than 230 bp (the maximum available from Agilent), randomly generated filler sequences were added evenly up- and downstream of the target ORF (keeping primers at the extremities) to maintain 230 bp final length.

Scripts (see data availability) were used with the following arguments:

python remove similar.py human denovo sorfs.csv

The resulting filtered file was used as input to generate a single subpool of oligos as follows:

python build oligos.py -i human denovo sorfs.unique.csv -s e coli -n 3750

-l 230 -r 1 -d primers.db -rf 1 -fL GGTCTCCA -fR GGCTCCCGAGACC

The two resulting .csv files containing separate oligo pools were concatenated and oligos ordered from Agilent.

### Age group classification

Ages of the *de novo* sORF proteins were assigned either based on sequence homology within outgroup genomes as described in above termed ”BLAST age” or based on transcription termed ”transcription age”. The age of each sORF protein corresponds to the hit in the furthest species from human on the evolutionary timescale. Age one corresponds to human specific *de novo* sORF proteins, two to sORFs present up to chimpanzee, three up to gorilla, four up to orang utan, five up to macaque and six up to mouse (see **Figure S10**). To determine the transription age, we used the locations of each sORF in the outgroup genomes to search the RNA-seq dataset from **wang°transcriptome°2020** covering brain, testis and liver tissues in human, macaque and mouse. We calculated TPM values for each sORF in each species-tissue combination by counting reads with HTseq (Anders et al. 2015). TPM values were calculated using the region defined by each BLAT hit in a given species. Where more than one BLAT hit was kept in a given species, the highest TPM value across all hits was kept.

### Prediction of protein properties

A number of sequence properties were predicted for the translated products of all library variants. Intrinsic structural disorder (ISD) was calculated using flDPnn (Hu et al. 2021) with a disorder threshold of 0.5 and mean, median, and percentage of disordered residues were extracted using multifasta fldpnn.py. Secondary structure content was predicted using NetSurfP 3.0 (Høie et al. 2022) and percentages for secondary structure elements extracted using netsurf predictions.py. For full structural prediction we used ESMFold (Lin et al. 2023). Based on the top ranked model we calculated N- to C-terminal distance, average solvent-accessible surface area (ASA) and secondary structure content using DSSP (Kabsch and Sander 1983), and radius of gyration (see script esm predictions.py).

### pETMF vector construction

To generate the FRET folding sensor we started with an in-house prepared pET09 backbone. pET09 is a modification of pET24a(+) (Milipore-Sigma, Burlington, MA) backbone with MCS switched for cloning cassette containing BsaI recognition sites. Plasmid harboring mVenus was obtained from Addgene (catalogue no.: 103986) and mTurquiose2 was a gift from Ondrej Havranek (coding sequence corresponding to Addgene catalogue no.: 61602). To make a FRET pair fusion cassette, the genes of fluorescent proteins (FPs) were furnished with cloning elements by PCR. BsmBI sites were added to 5’ end of mTurquiose2 and 3’ end of mVenus sequence creating matching overhangs to the BsaI sites in pET09 backbone. The opposite termini of FPs were rigged with: (i.) Eco53kI recognition sites and GGS motive to link the insert with the FPs; (ii.) BsaI sites to be used for cloning of the libraries; (iii.) BsmBI sites to assemble the FRET cloning cassette. pET09 vector was opened with BsaI enzyme and dephosphorylated with rSAP enzyme (NEB, Ipswich, MA), PCR amplicons of furnished FPs were digested with BsmBI enzymes and following clean-up all three elements were ligated with T4 DNA ligase (NEB, Ipswich, MA).

### Construction of controls

We generated plasmids for single expression of donor/acceptor and fusion expression of the FRET pair separated by either glycine/serine linkers of different lengths, 2 characterised random proteins or 6 well characterised *E.coli* proteins with diverse length and properties. In the case of GS controls, we used PCR with long protruding primers to add GGS sequences and BsaI sites directly to the genes of mTurquiose2 and mVenus and assembled them with the pET09 via NEBridge^®^ Golden Gate Assembly Kit (BsaI-HF^®^ v2) (NEB, Ipswich, MA). Control proteins were PCR amplified from either *E.coli* DH5 αgenome or in-house generated constructs, to contain BsaI sites compatible with pETMF vector and subcloned via GGA. *E. cloni* 10G strain (LGC, Biosearch Technologies, Hoddesdon, UK) was used for all the cloning steps and plasmid amplification. Once confirmed by Sanger sequencing, constructs were retransformed into *E.coli* BL21 (DE3) for expression and cultures were stored as glycerol stocks at -80 °C.

### Library synthesis and cloning

To obtain oligo pools of the libraries, we used Agilent’s SurePrint Oligonucleotide Library Synthesis (OLS). Individual libraries were PCR amplified from the pool to contain BsaI sites compatible with pETMF vector. Purified PCR product and vector were added to 20 µl Golden Gate assembly reaction in 2:1 molar ratio and run overnight in 5-minute cycles of 16 °C and 37 °C. The reaction mix was purified with Monarch^®^ PCR & DNA Cleanup Kit (NEB, Ipswich, MA). Next, 1 µl of purified reaction mix was electroporated to 25 µl of in-house prepared *E. cloni* 10G cells and incubated overnight at 37 °C on a LB-agar plate supplemented with 50 µg/ml kanamycin (same for all following growth media). The colonies were pooled and plasmid DNA isolated with Zyppy Plasmid Miniprep Kit (ZymoGenetics, Inc, Seattle, WA). 50 ng of pETMF carrying either of the libraries was electroporated to 25 µl of *E.coli* BL21 and incubated overnight at 30 °C on a LB-agar plates containing kanamycin. The plates were washed with cold PBS containing kanamycin to collect the colonies. Finally, the mixture of scraped cells was used for both plasmid DNA isolation (serving as “presort” sample in NGS analysis) and to prepare 1 ml glycerol stocks (20% v/v) with OD600 adjusted to 1.

### Protein expression

Cells carrying control protein plasmids were inoculated from glycerol stocks, grown overnight at 37 °C, reinoculated to fresh media and grown at 37 °C. After reaching OD600=0.6, cells were cooled down and expression induced with IPTG to 0.5 mM concentration followed by expression at 25 °C for 16 hours. Glycerol stocks of BL21 cells carrying library DNA were thawed on ice, diluted with fresh LB with kanamycin to OD600=0.2 and grown shaking at 37 °C until reaching cell density 0.6 (approximately 50 minutes). Cells were cooled down, IPTG added to 0.5 mM and expression carried out at 25 °C for 16 hours.

### Measurement of donor fluorescence lifetime

Expressing cells were harvested (5 min, 4000 x g, 4 °C) and washed 3 times with cold PBS. Density of the cultures was adjusted to approximately 1, while keeping the cells on ice. Fluorescence measurements were carried out on Photoluminescence Spectrometer FLS 1000 (Edinburgh Instruments Ltd., Livingston, UK). We used Xenon lamp for steady-state measurements to obtain emission spectra of mTurquoise2. For time-resolved measurements a 405 nm picosecond diode laser with repetition rate set to 10 MHz was used to collect a fluorescence decay at 473 nm emission in three technical replicates.

### Flow cytometry

Following the overnight expression, the OD600 of cell cultures was adjusted to 1. The cells were collected (5 min, 4000 x g, 4 °C), washed three times and diluted 30 times in cold, filtered PBS. The cytometer BD FACSAria Fusion was equipped with a 70 µm nozzle and ND (neutral density) 1.0 filter. The sample chamber temperature was set to 4 °C and in the case of sorting, the collection tube temperature was set to 30 °C. We used the 405 nm violet laser with 450/50 bandpass filter for donor emission, 525/50 filter for FRET emission and 488 nm blue laser with 530/30 filter for acceptor detection. SSC detection threshold was set to 300 to capture the size of *E. coli* cells. After initial gating for size and shape (SSC-A x FSC-A and FSC-H x FSC-A), we used donor versus acceptor channel plot to gate population positive for both fluorophores (P1). To set the gates for FRET positive/negative sorting, the P1 population was projected as a histogram of a parameter derived from the ratio of the FRET channel and the donor channel (FRET ratio). The cells were sorted to rich recovery medium (LGC, Biosearch Technologies, Hoddesdon, UK) in a 1.5 ml centrifugation tube. Following the sorting, the cells were plated on a LB-agar+kanamycin plate and incubated at 30 °C for 16 h. Finally, the colonies were scraped to LB+kanamycin and used either for subsequent round of expression and sorting or the plasmid DNA was extracted. Stocks from all rounds of sorting were then cultured and expressed in a single experiment and recorded on BD LSRFortessa cytometer.

### High-throughput sequencing

The plasmid pools recovered from the FACS experiments were used to generate PCR amplicons for subsequent NGS analysis. We used Q5^®^ High-Fidelity DNA Polymerase (NEB, Ipswich, MA) with 50 ng of plasmid DNA per 50 µL of PCR as a template and the reaction was run for eleven cycles. Primers were designed to anneal to the pETMF backbone and introduce nine sets of barcodes (see Zenodo). Following a clean-up, the amplicon size distribution for selected samples was obtained by the Agilent 2100 Bioanalyzer (Agilent Technologies, Inc., Santa Clara, CA). In total, four sequencing libraries were generated by NEBNext^®^ Ultra ™II DNA Library Prep Kit for Illumina ^®^ (NEB, Ipswich, MA). Sizes and concentrations of the sequencing libraries were again verified by the Agilent 2100 Bioanalyzer. Finally, the samples were pooled as a part of a larger run on Illumina NextSeq platform. NextSeq™1000/2000 P1 Reagents with 600 cycles were used and we dedicated 340-440K reads per sample. Reads were merged, trimmed and filtered to remove low quality reads using the fastp suite (S. Chen et al. 2018). Reads were mapped to CDS sequences of library DN and R, respectively, using the Burrows-Wheeler Alignment (BWA) MEM algorithm (Li and Durbin 2009). SAMtools was used for conversion to SAM file format, sorting and indexing (Li, Handsaker, et al. 2009). Reads mapped to each variant were then counted using the HTSeq python module (Anders et al. 2015) using the script sam counter.py.

### Enrichment analysis

Library sequences were filtered out before enrichment analysis if the total read count number across replicates was below 50 and if the sequence was not present in at least two replicates including the presorted library. Enrichment of single sequences was calculated using the python implementation of DESeq2 (Love et al. 2014) comparing sorted samples after round one and two to the presorted samples. For analysis only the positive log fold changes with a significant adjusted p-value below 0.05 were used. To check for statistical differences we used SciPy (Virtanen et al. 2020). To analyse differences between the FRET-positive and FRET-negative groups we applied standard t-test, for differences between multiple groups we used Kruskal-Wallis test with Dunn post-hoc test. To check whether the numbers of sequences sorted into FRET-positive or FRET-negative were non-random between age groups we applied Chi2 test.

### Statistical modelling

All statistical modeling was done using R version 4.3.1 (Ihaka and Gentleman 1996). The data was split first into *de novo* and random sequences and then into positive and negative FRET data sets. These data sets were further divided randomly in half into a training and test data set. Models were then created using the glmnet package (Tay et al. 2023) on the respective training data set. Cross-validation was run with ten folds using type-measure deviance. This was done with four different alpha values; 1, 0.7, 0.5 and 0.3. The chosen model was the minimum lambda model with the lowest deviance out of the 4 alpha values. Predictive plots were then created using a custom link function from the coefficients of the training set models and plotted against the test data. For random sequences, the model intercept was included while for the *de novo* sequences, it was not.

### Data availability

Code used for library design, prediction of protein properties and analysis is available at: https://zivgitlab.uni-muenster.de/ag-ebb/de-novo/fret-facs/

All used protein sequences, result files and raw NGS reads are available at Zenodo under: https://zenodo.org/doi/10.5281/zenodo.10498066

## Acknowledgements

M.A., F.B., E.B.B. and K.H. received funding from Volkswagen foundation grant code 98183. M.A. and E.B.B.received funding from HFSP grant HFSP - RGP004/2023. F.B. and K.H. received funding from Primus grant PRIMUS/20/SCI/012 from Charles University. The project received funding from the EU under the Horizon 2020 Research and Innovation Framework Programme No. 722610 and the Erasmus+ programme. We thank Tereza Neuwirthova for running ESMFold predictions using computational resources provided by the e-INFRA CZ project (ID:90254), supported by the Ministry of Education, Youth and Sports of the Czech Republic. Gustavo Fuertes Vives and Tatsiana Charnavets for their help with the fluorescence measurements and Ondrej Havranek for kindly providing the mTurquiose2 construct and discussing the flow cytometry results. We acknowledge the Imaging Methods Core Facility at BIOCEV for their support with obtaining flow cytometry data presented in this paper. We thank Max Gantz and Lars Eicholt for discussion of the results. Figure 1 was created with BioRender.com.

---

**Figure S1:**
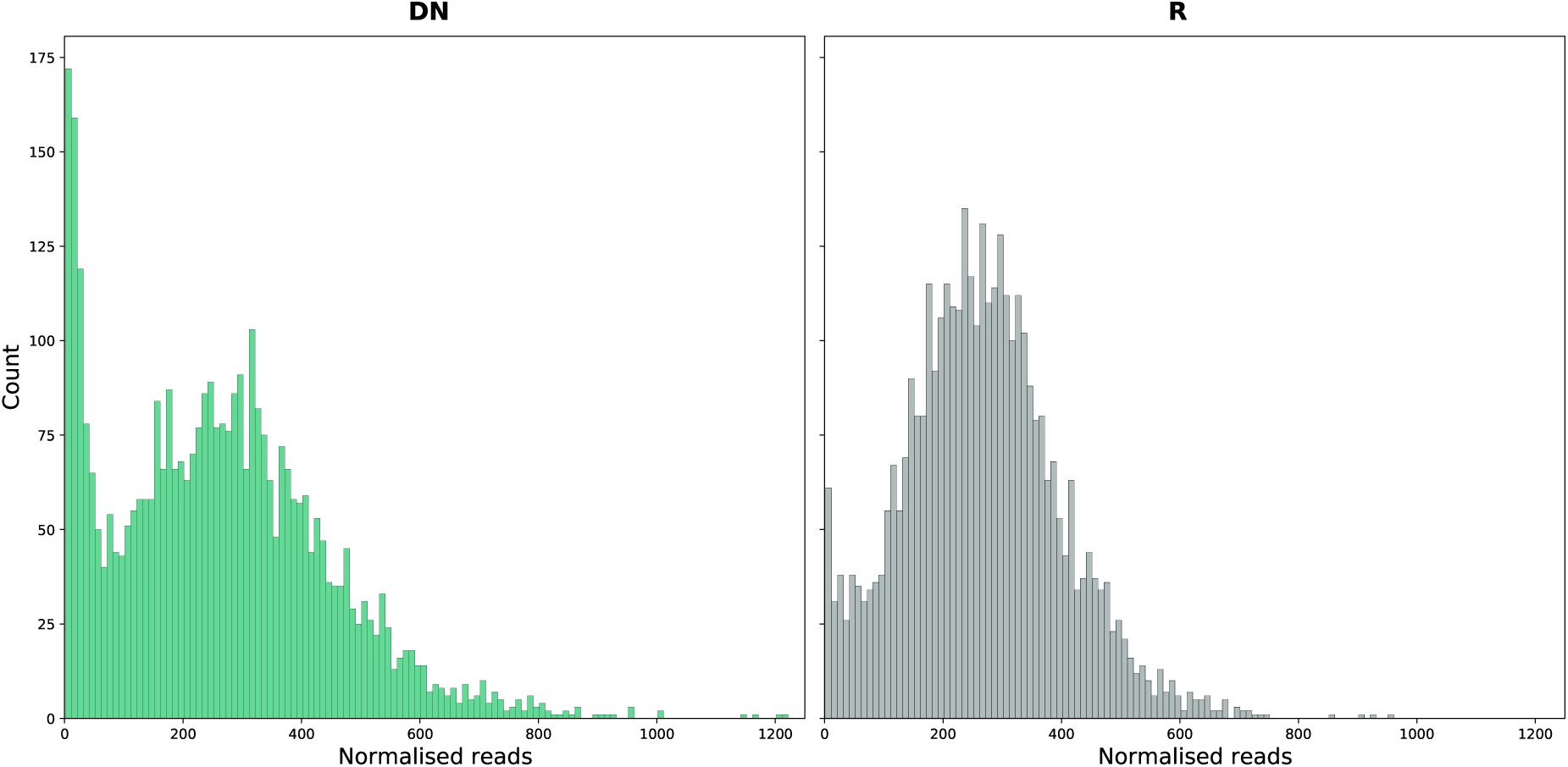
Depth-normalised read count distributions for library DN and R cell-free templates.

**Figure S2:**
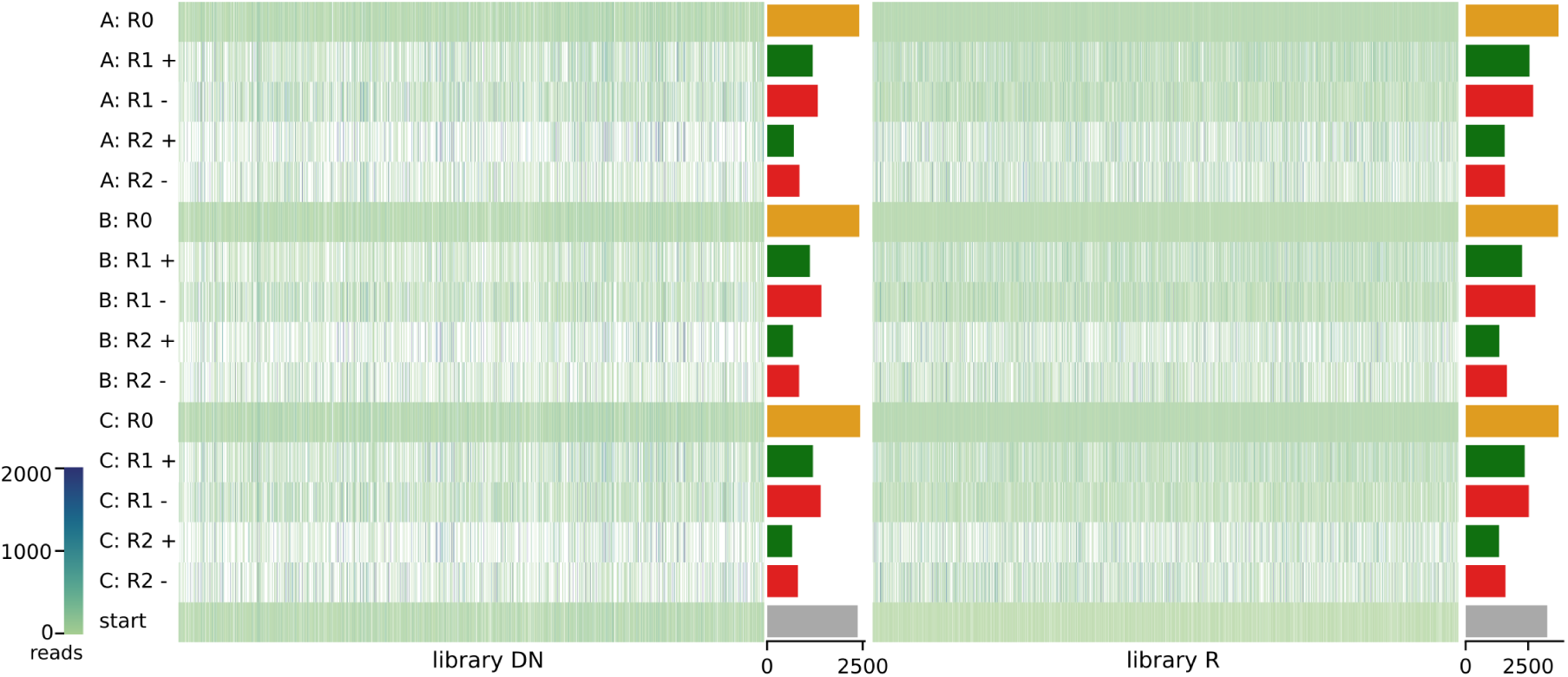
Read counts per sequence for libraries DN and R for presort (0), and both rounds of sorting (1-2) into FRET-positive (+) and FRET-negative (-) for all replicates (A-C). Number of unique sequences are shown in bar plots on the right.

**Figure S3:**
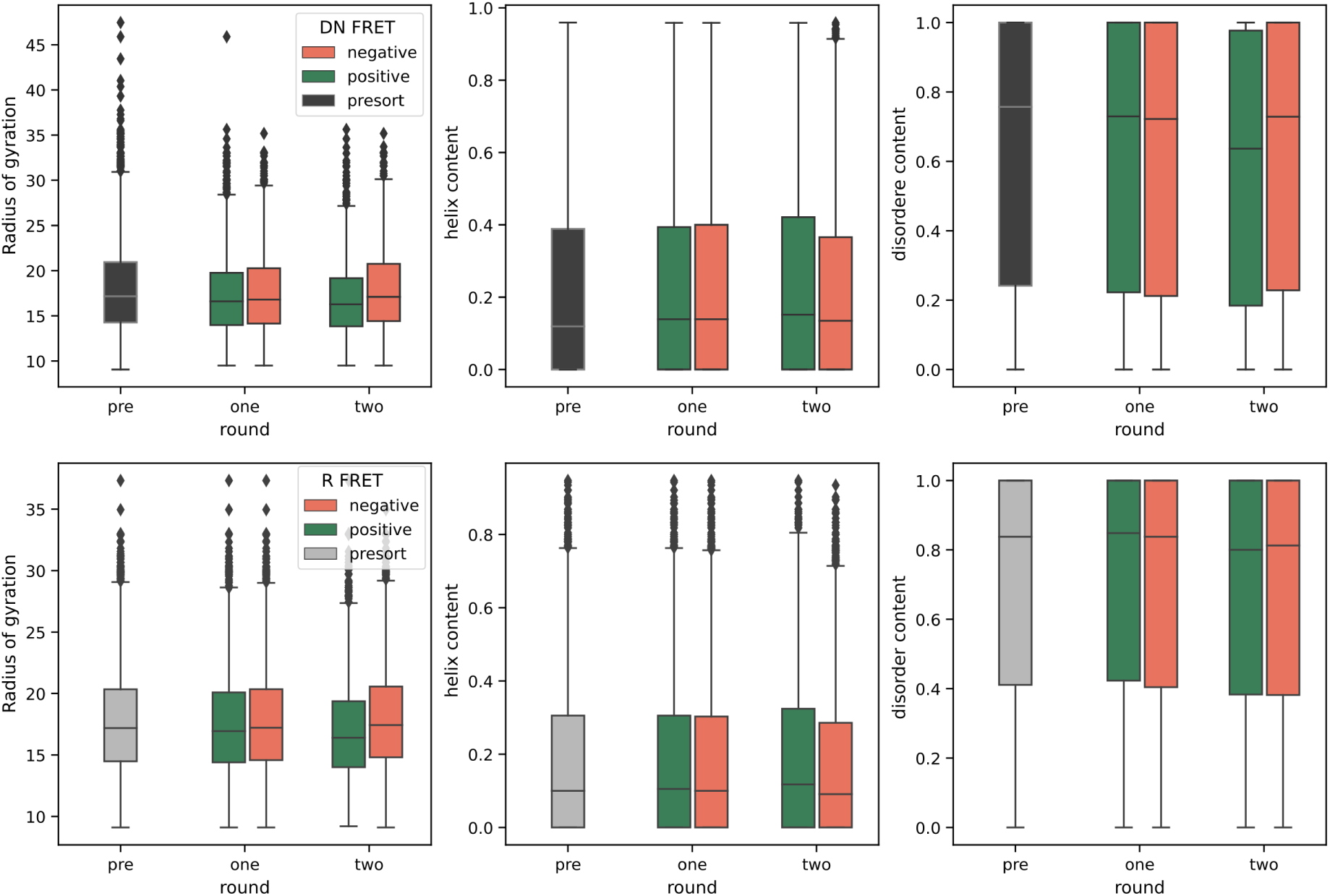
Radius of gyration, α-helix content and disorder content of all unique sequences in Presort, positive and negative samples across rounds for libraries DN and R.

**Figure S4:**
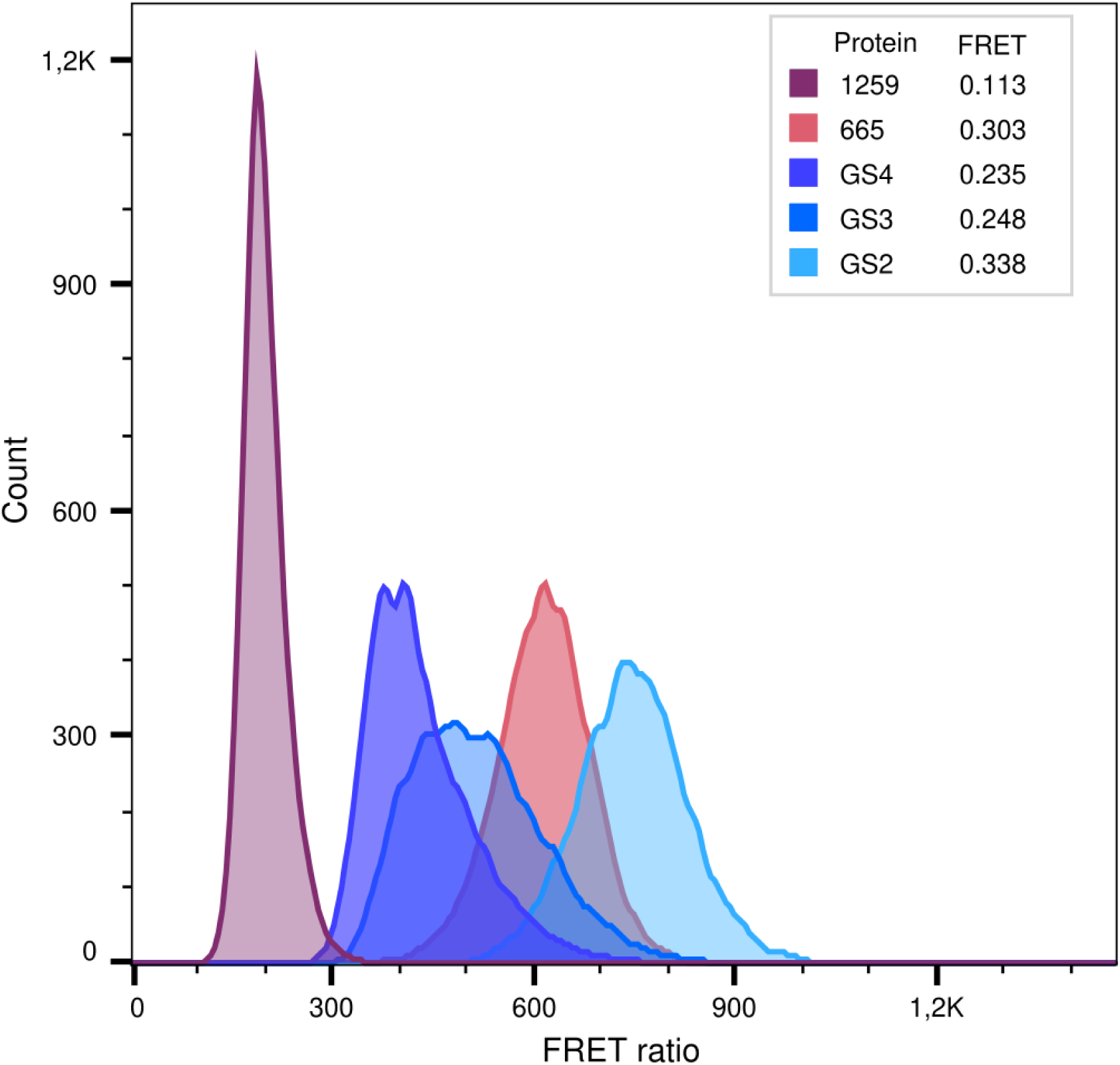
FRET ratio parameter from cytometric analysis of selected control proteins with corresponding FRET efficiencies from fluorescence lifetime measurements.

**Figure S5:**
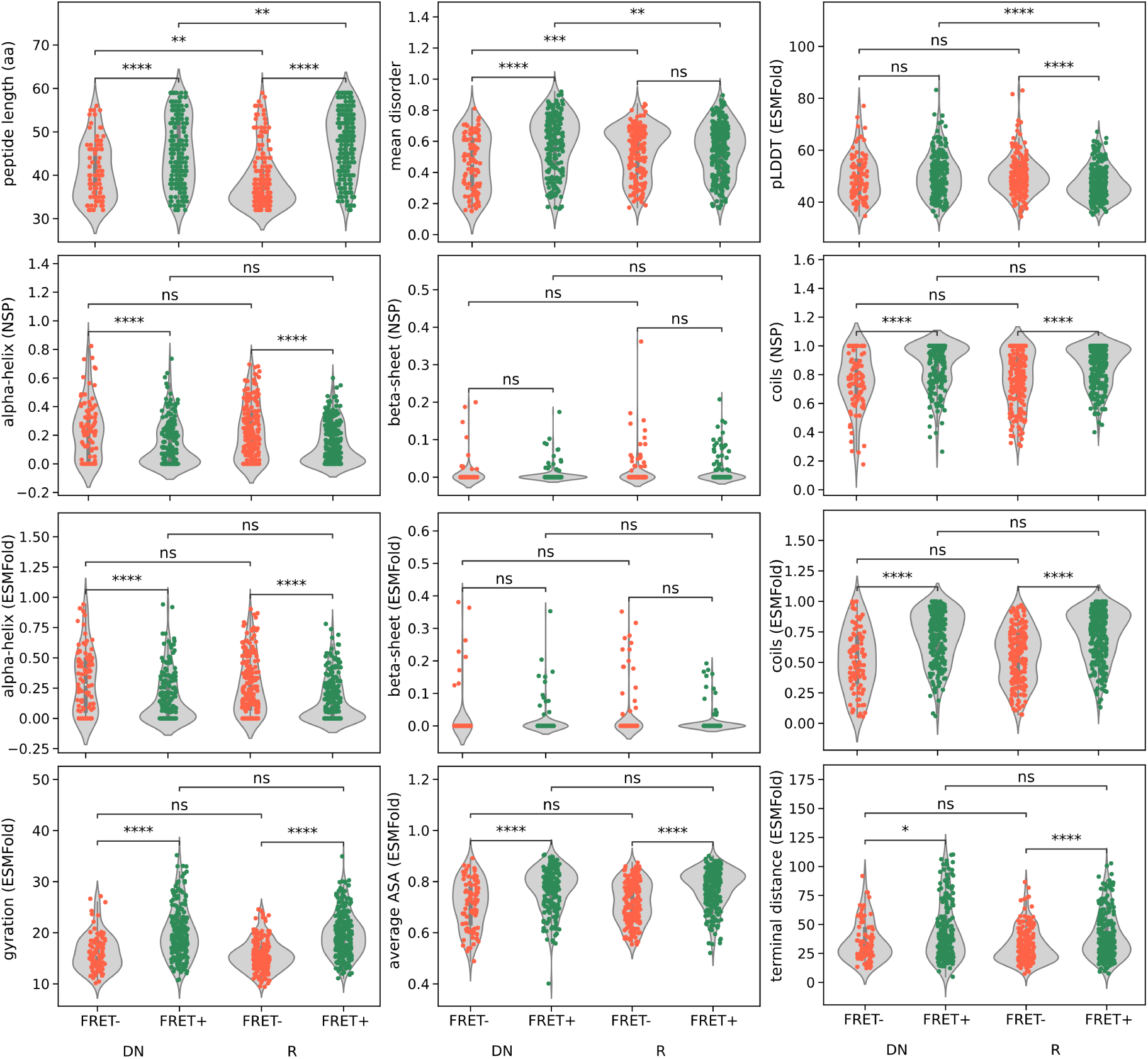
FRET-positive (green) and FRET-negative (red) enriched sequences from pre-sort to round two of sorting for libraries DN and R. Stars indicate significance calculated with post-hoc Dunn test after Kruskal-Wallis test (p-value *<* 0.05 * *<* 0.01 ** *<* 0.001 *** *<* 0.0001 ****

**Figure S6:**
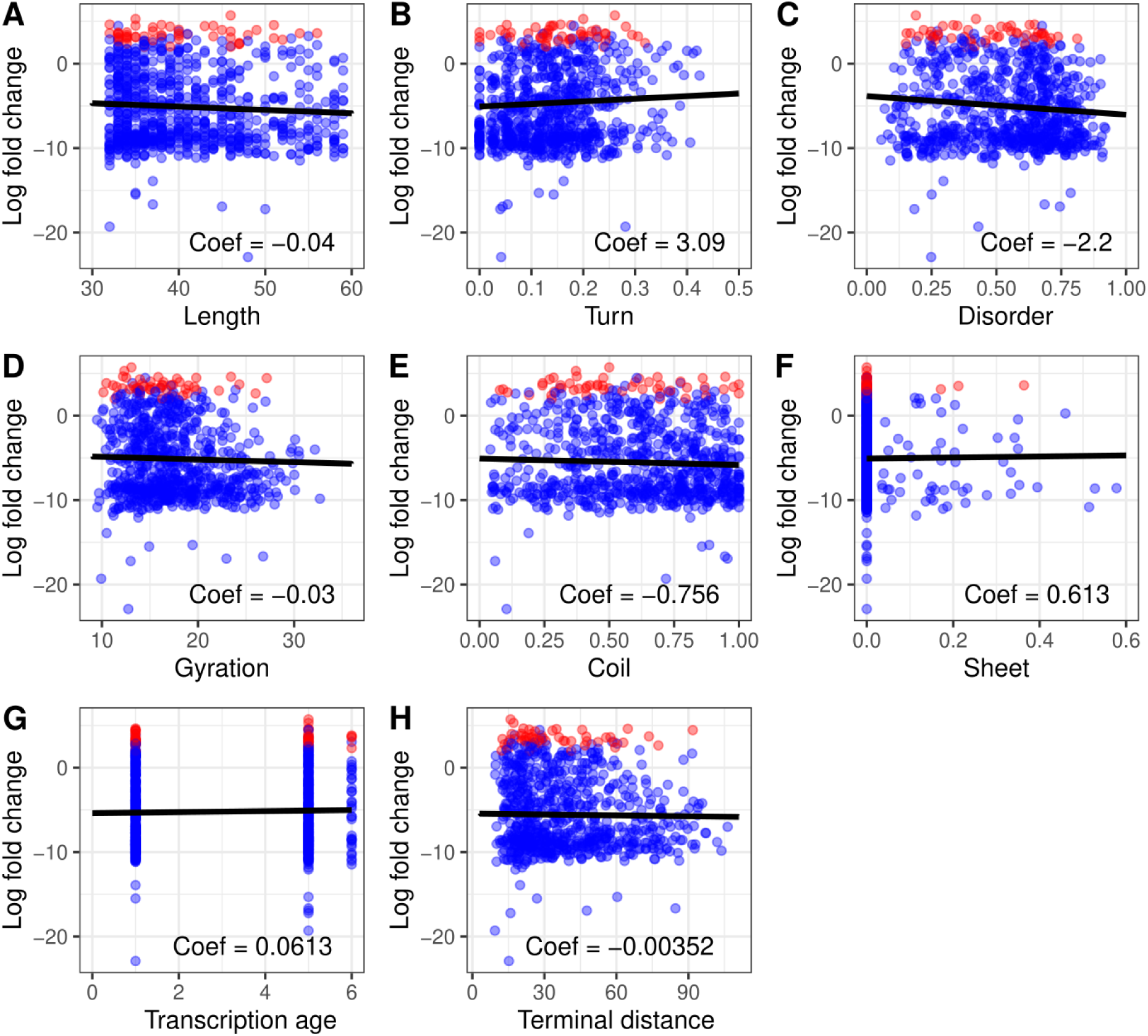
Predictive plots for FRET-positive *de novo* sORF proteins enriched from pre-sort to round two. The points are the test data set with significantly enriched sequences in red. The line is the coefficient of the particular predictor with all other variables either set to zero or to the median value. Note that in an an elastic net type regression as used here, the uncertainty is not calculable in the same way as a normal regression, so no confidence intervals are added.

**Figure S7:**
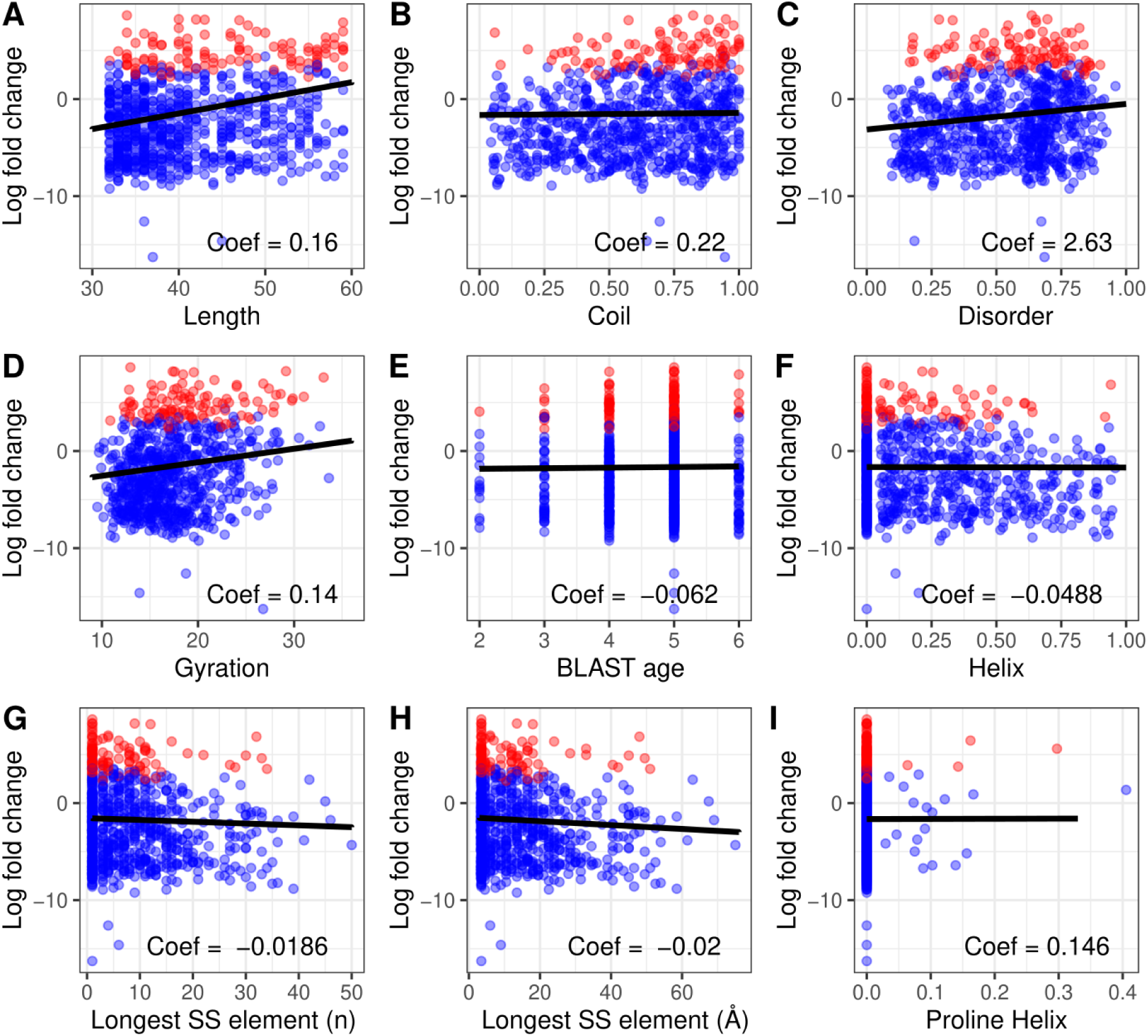
Predictive plots for FRET-negative *de novo* sORF proteins enriched from pre-sort to round two. The points are the test data set with significantly enriched sequences in red. The line is the coefficient of the particular predictor with all other variables either set to zero or to the median value. Note that in an an elastic net type regression as used here, the uncertainty is not calculable in the same way as a normal regression, so no confidence intervals are added.

**Figure S8:**
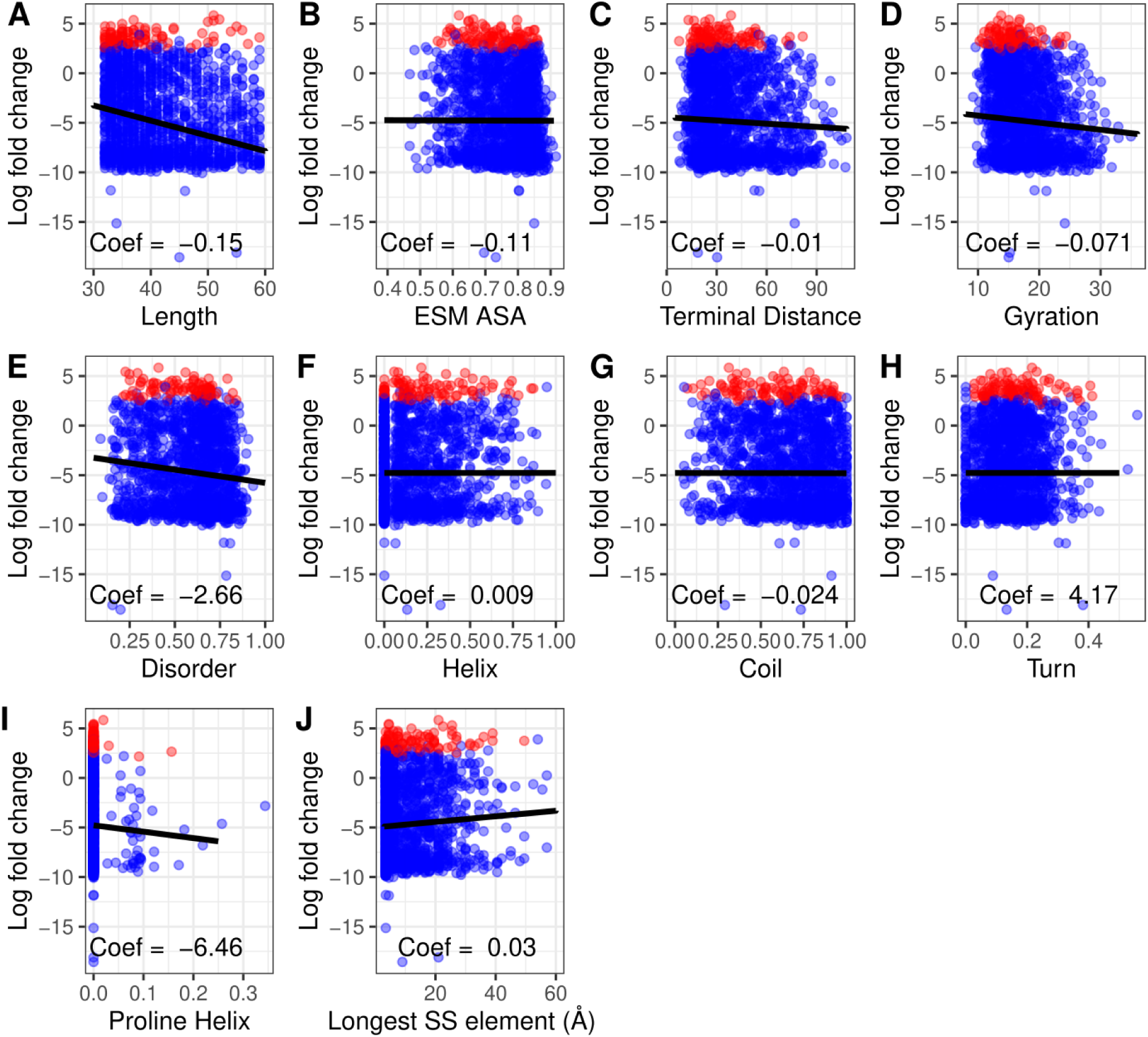
Predictive plots for FRET-positive randomly generated sORF proteins enriched from presort to round two. The points are the test data set with significantly enriched sequences in red. The line is the coefficient of the particular predictor with all other variables either set to zero or to the median value. Note that in an an elastic net type regression as used here, the uncertainty is not calculable in the same way as a normal regression, so no confidence intervals are added.

**Figure S9:**
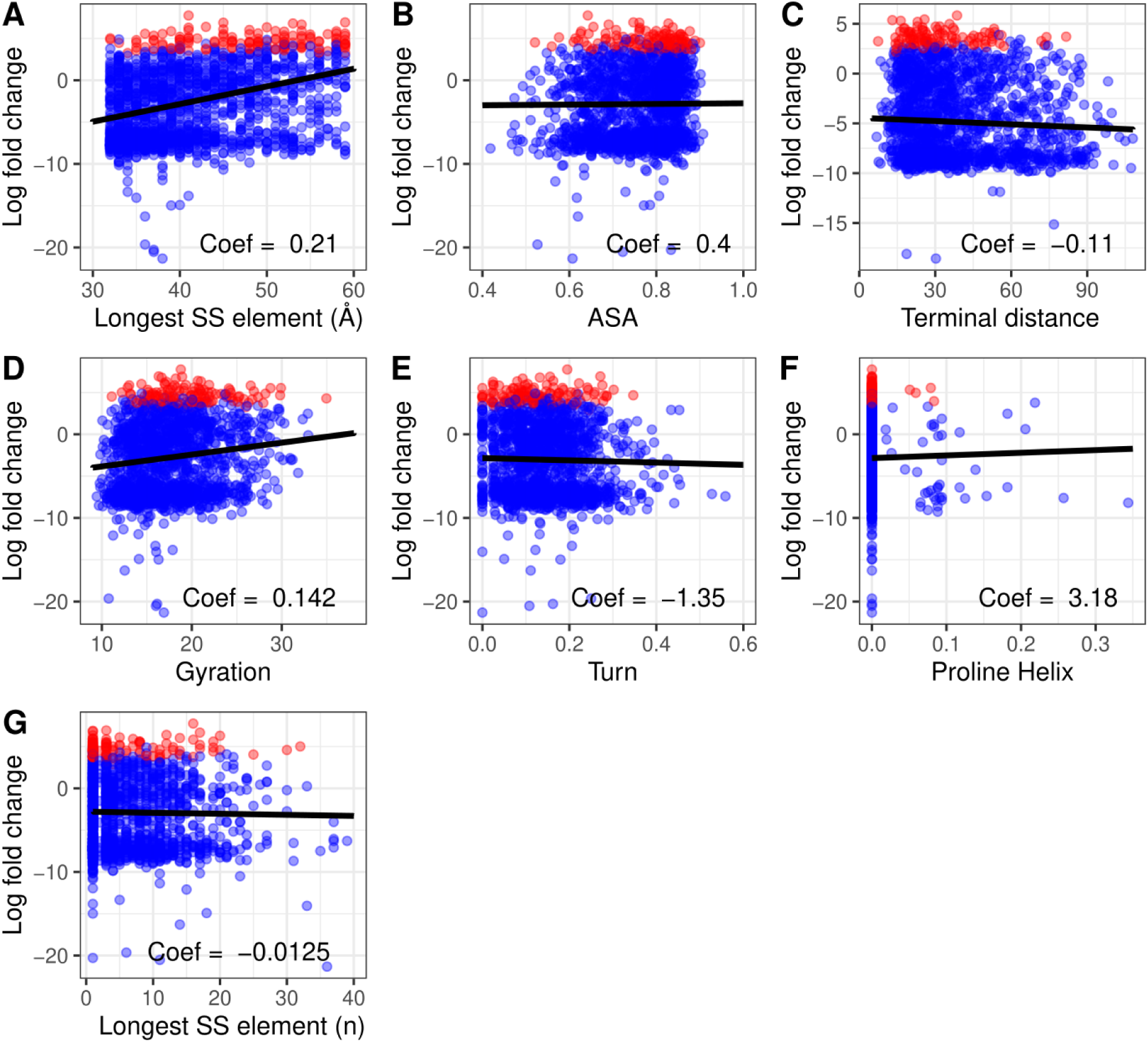
Predictive plots for FRET-negative randomly generated sORF proteins enriched from presort to round two. The points are the test data set with significantly enriched sequences in red. The line is the coefficient of the particular predictor with all other variables either set to zero or to the median value. Note that in an an elastic net type regression as used here, the uncertainty is not calculable in the same way as a normal regression, so no confidence intervals are added.

**Figure S10:**
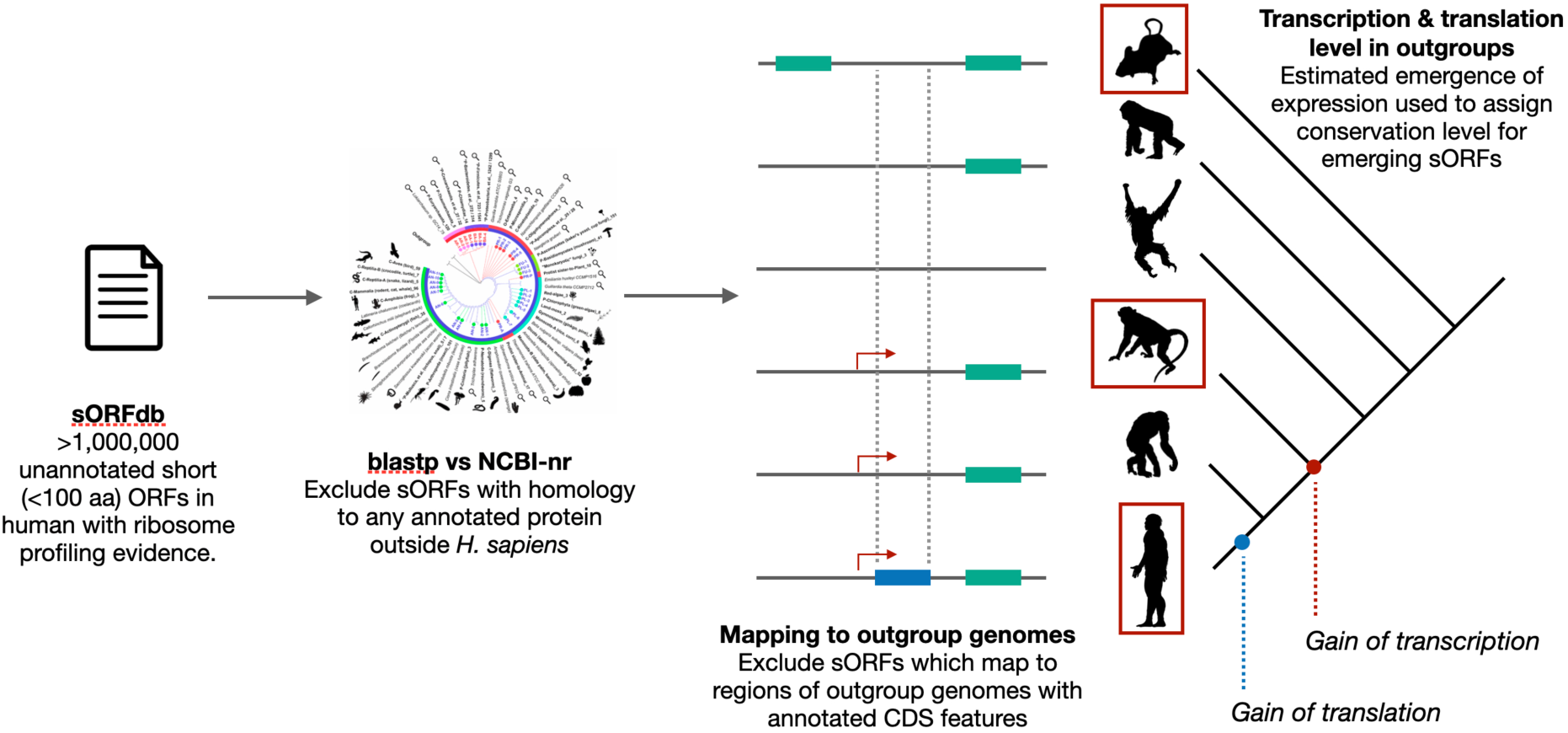
Identifying a pool of putatively *de novo*-emerged human-specific sORFs.

**Table S1:**
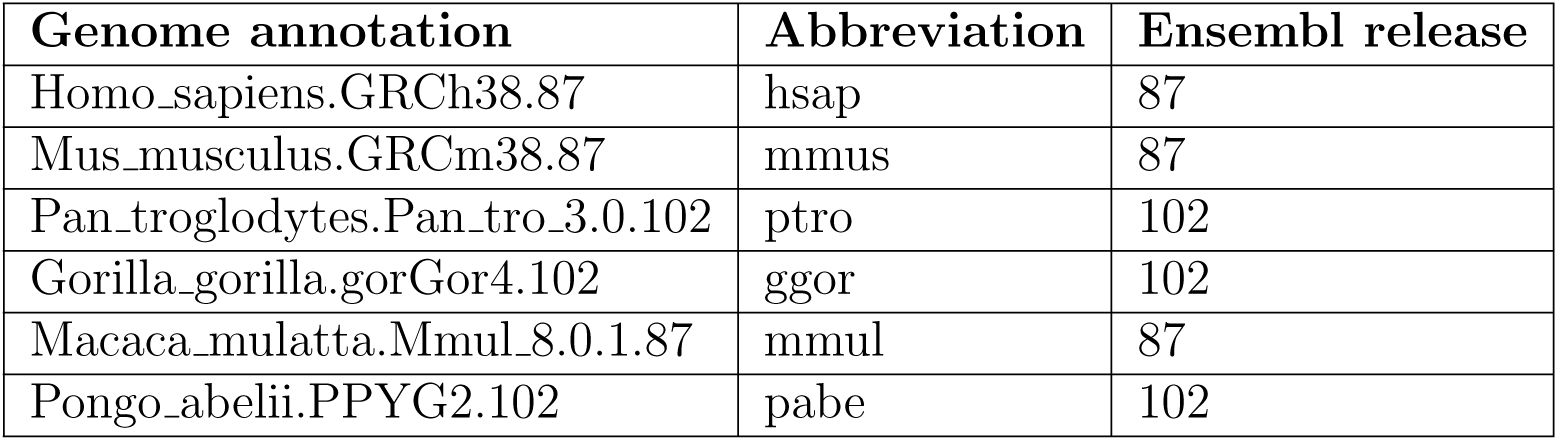
Genome annotations used for BLAT mapping of human sORFs.

